# tRNA synthetase activity is required for stress granule and P-body assembly

**DOI:** 10.1101/2025.03.10.642431

**Authors:** Max Baymiller, Noah S. Helton, Benjamin Dodd, Stephanie L. Moon

## Abstract

In response to stress, translation initiation is suppressed and ribosome runoff via translation elongation drives mRNA assembly into ribonucleoprotein (RNP) granules including stress granules and P-bodies. Defects in translation elongation activate the integrated stress response. If and how stalled ribosomes are removed from mRNAs during translation elongation stress to drive RNP granule assembly is not clear. We demonstrate the integrated stress response is induced upon tRNA synthetase inhibition in part via ribosome collision sensing. However, saturating levels of tRNA synthetase inhibitors do not induce stress granules or P-bodies and prevent RNP granule assembly upon exogenous stress. The loss of tRNA synthetase activity causes persistent ribosome stalls that can be released with puromycin but are not rescued by ribosome-associated quality control pathways. Therefore, tRNA synthetase activity is required for ribosomes to run off mRNAs during stress to scaffold cytoplasmic RNP granules. Our findings suggest ribosome stalls can persist in human cells and uniquely uncouple ribonucleoprotein condensate assembly from the integrated stress response.

## Introduction

The integrated stress response (ISR) is a conserved signaling pathway in which phosphorylation of the translation initiation factor eIF2α causes translation inhibition and gene expression reprogramming (*1*, *2*). During the ISR, inhibition of translation initiation and ribosome runoff release mRNAs from the translating pool. Ribosome-free mRNAs make multivalent interactions with RNA-binding proteins and other RNAs, and together with protein-protein interactions drive the assembly of biomolecular condensates called stress granules (SGs) and processing bodies (P-bodies or PBs) (*3–6*). These stress-induced ribonucleoprotein (RNP) granules can sequester RNAs and RNA-binding proteins away from the cytoplasm (*7–13*) and are enriched in translationally repressed mRNAs (*14–21*). SGs and PBs have been implicated in the cellular resilience to stress and disease contexts (*22–26*), yet are also associated with a maladaptive or cytotoxic stress response (*27–30*). The functions of RNP granules in cellular resilience to stress and the ISR may be context-dependent and are still being resolved.

Stressors such as impaired tRNA charging, amino acid deprivation, UV, and other RNA damaging agents inhibit translation elongation and activate the ISR through GCN2 (*31–38*)). GCN2 can be activated *in vitro* by binding uncharged tRNAs (*39–41*) and/or the ribosomal P-stalk (*42*, *43*). GCN2 activation is associated with collisions between elongating and stalled ribosomes that occur when mRNAs are damaged, amino acids are depleted, or upon genetic depletion of tRNA and ribosome rescue factors (*32*, *33*, *38*, *39*, *42*, *44–47*). Recognition of ribosome collisions by GCN2 may be mediated by interactions with the ribosomal P-stalk and/or by the cofactor GCN1 that has been structurally placed at the disome interface (*46*, *47*). In these contexts, GCN2 activation mediates a negative feedback loop between translation elongation and initiation to inhibit further ribosome stalling events and reduce the production of partially generated proteins that can misfold and aggregate. The interplay between translation elongation and initiation through GCN2 suggests that this feedback loop is critical for cellular adaptation to stress (*33*, *38*, *39*, *42*, *44–47*).

It is not clear whether and how RNP granules assemble during stress conditions that activate the ISR and inhibit translation elongation. Ribosome-free mRNA is essential to drive and scaffold canonical stress granule assembly in wild-type cells that do not exogenously over-express core SG components. Chemical inhibitors of elongation such as cycloheximide trap ribosomes on mRNAs and prevent SG and P-body formation (*11–13*, *48–51*). Ribosome-bound mRNAs are not stably sequestered within SGs (*7*, *52*), and mRNA association with as few as one ribosome blocks SG formation (*48*), indicating ribosome occupancy and mRNA condensation into SGs are incompatible (*4*). It is thus anticipated that RNP granule assembly will be limited during stresses that cause ribosome stalling. However, multiple mechanisms exist to remove stalled ribosomes from mRNAs, including ribosome-associated quality control (RQC) (*53*, *54*) and ribosome rescue factors such as HBS1/PELO (*55*, *56*) and GTPBP1/2 (*57*, *58*). Yet, these pathways can be overwhelmed (*44*, *59*) and emerging evidence suggests they cannot resolve some types of ribosome collisions (*32*, *60*, *61*). It is unknown whether RQC/ribosome rescue pathways or other mechanisms enable RNP granule assembly during translation elongation stress.

Inhibitors of tRNA charging provide an opportunity to dissect the mechanistic links between ribosome stalling, RNP granules, and the ISR. Halofuginone is a potent and specific inhibitor of the prolyl-tRNA charging activity of glutamyl-prolyl tRNA synthetase (EPRS1) (*36*). Halofuginone activates the ISR via GCN2 (*36–38*, *62–64*) and increases ribosome occupancy at proline codons (*62*, *64*) suggestive of ribosome stalling. Activation of GCN2 by halofuginone has been suggested to occur via direct sensing of uncharged tRNA without a requirement for ribosome collisions (*38*). However, ribosome collisions occur after genetic depletion of tRNA synthetases (*65*, *66*) and tRNA (*45*, *58*) in association with GCN2 activation. Further, a prior study found TIA-1/TIAR+ ‘riboclusters’ reminiscent of SGs are induced upon low-level tRNA synthetase inhibition (*67*), in contrast to an earlier study that reported SGs do not form upon treatment with the threonyl-tRNA synthetase inhibitor borrelidin (*51*). Halofuginone and other tRNA synthetase inhibitors have therapeutic potential for a range of disease contexts including cancers, fibrosis, autoimmunity, parasitic infections, and diabetes (*37*, *68–71*). Mutations in tRNA synthetases are also associated with heritable peripheral neuropathies and multisystem disorders (*72–79*), including loss-of-function EPRS1 variants (*80–82*). Further, ISR activation has been observed in affected motor neurons in animal models of tRNA synthetase disease (*83*, *84*). Therefore, elucidating how the ISR functions when tRNA charging is impaired may suggest insights into mechanisms of the stress response, disease, and therapeutic modes of action.

In this study, we investigated the impact of tRNA synthetase inhibitors on translation elongation, the ISR, and stress-induced RNP granules. We found that tRNA synthetase inhibitors did not induce SG assembly despite robustly activating the ISR to levels equal to stresses that cause SG formation. Furthermore, tRNA synthetase inhibitors prevented SG formation during other stresses. Loss of tRNA charging also disassembled P-bodies and prevented them from being induced by other stresses. Impaired RNP granule assembly upon tRNA synthetase inhibition was associated with global translation suppression and retention of mRNAs in polysomes, suggesting global ribosome stalling. We found that the RQC pathway initiated by the ubiquitin ligase ZNF598, which recognizes collided ribosomes, was activated at a low level upon tRNA synthetase inhibition and contributed to the ISR, but failed to resolve stalled ribosomes. The peptidyl-tRNA mimic puromycin freed mRNAs from polysomes and rescued RNP granule assembly during halofuginone stress. Surprisingly, halofuginone-induced ribosome stalls persisted for over 16 hours. Together, these results suggest that stresses that inhibit translation elongation can uncouple RNP granule assembly from the ISR due to unresolved, persistent ribosome stalls. These persistent stalls are the result of the limited cellular capacity to detect elongation defects via ribosome collisions or uncollided, stalled ribosomes.

## Results

### Inhibition of tRNA charging activates the integrated stress response (ISR) and suppresses translation

We first assessed whether acute inhibition of tRNA synthetases activates the ISR. We treated U-2 OS cells with micromolar levels of halofuginone to rapidly and completely inhibit prolyl-tRNA charging for 1 hour and assessed eIF2α phosphorylation and translation activity. We compared P-eIF2α levels upon halofuginone treatment to arsenite and thapsigargin stress, which activate the ISR via oxidative or endoplasmic reticulum stress (respectively). Consistent with prior studies (*36–38*, *62–64*), we observed that halofuginone significantly increased P-eIF2α levels (by 16.0 +/- 1.5 fold), which was comparable to P-eIF2α increases induced by arsenite (21.0 +/- 3.5 fold) and thapsigargin (13.8 +/- 2.6 fold) (**Fig. 1A**). We then applied bioorthogonal noncanonical amino acid tagging (BONCAT) (*85*) to measure bulk protein synthesis during stress. We found that halofuginone caused an 84.0 +/- 4.5% reduction in protein synthesis compared to unstressed cells as measured by BONCAT, which was comparable to the degree of translational suppression observed in arsenite and thapsigargin stresses (**Fig. 1B**). Therefore, inhibition of prolyl-tRNA charging activates the ISR and suppresses translation to a similar degree as canonical stressors.

**Figure 1.**
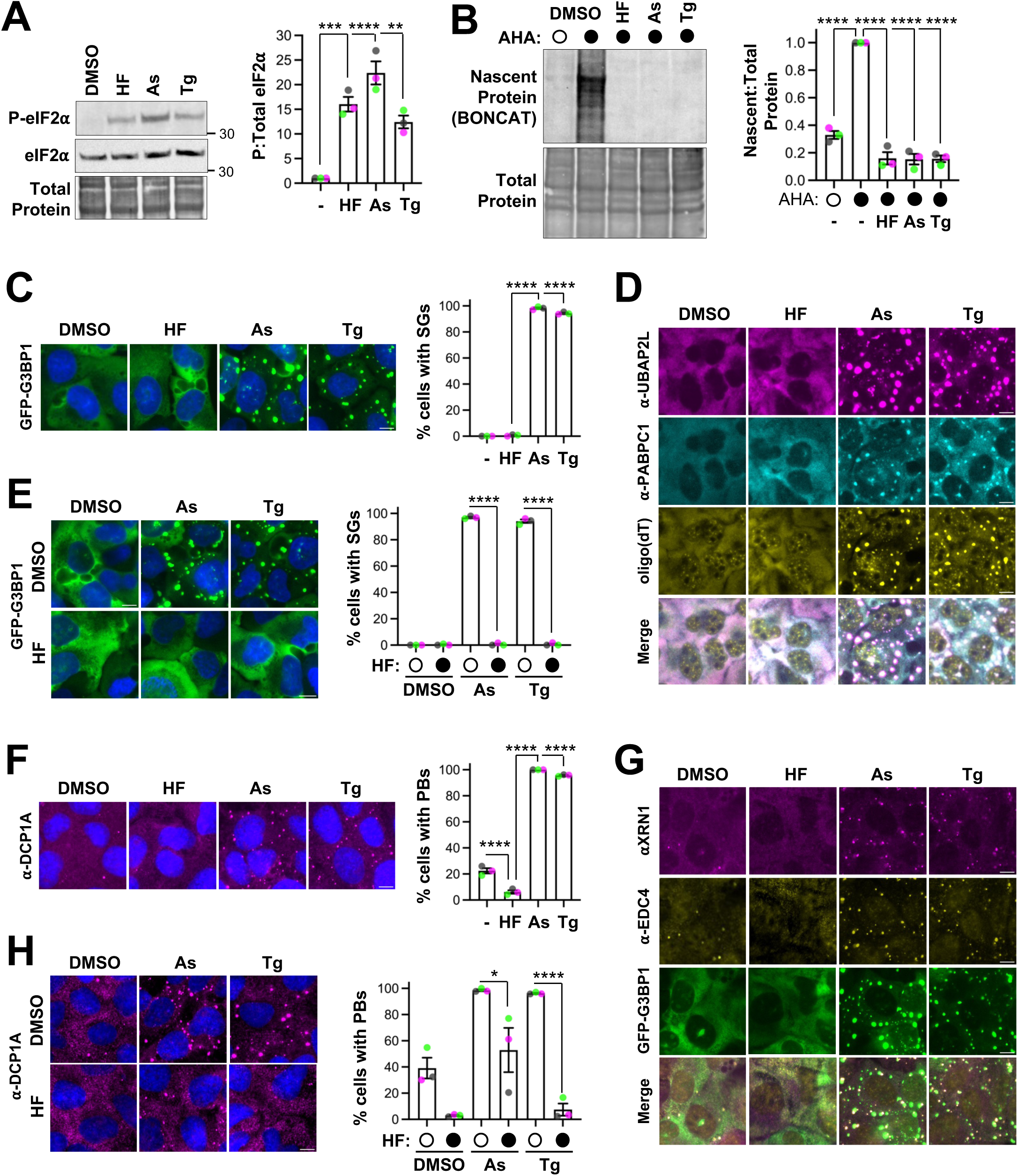
Inhibition of tRNA charging activates the ISR without inducing RNP granules. A. U-2 OS cells were treated with DMSO carrier (0.2%), halofuginone (HF, 20 µM), sodium arsenite (As, 250 µM), or thapsigargin (Tg, 1.25 µM) for 1 hour and western blotting for phosphorylated and total eIF2α done with total protein shown (n=3 independent replicates). Molecular weights (kDa) are shown on each blot. B. Translation protein is shown from n=3 independent replicates. C. U-2 OS cells stably expressing GFP-G3BP1 (green) were treated as in (A), fixed and nuclei stained with Hoechst (blue), and the percent cells with SGs was quantified from n=3 independent experiments (>305 cells counted in each condition). Scale bars: 10 µm. D. Cells were treated as in (C), and immunofluorescence and fluorescence *in situ* hybridization done to detect UBAP2L (magenta), PABPC1 (cyan), and poly(A) RNA with oligo(dT) probes (yellow) (n=2 independent experiments). E. Cells were treated with DMSO (0.2%) or 20 µM HF and DMSO (0.2%), 250 µM arsenite, or 1.25 µM thapsigargin for 1 hour, and the percent cells with SGs quantified (n=3 independent experiments with ≥287 cells counted per treatment). F. Cells were treated and immunofluorescence microscopy done as in (D) to detect P body marker DCP1A. Quantification of the percent cells with P bodies (PBs) is shown from n=3 independent experiments with ≥364 cells counted per treatment. G. Cells were treated as in (C) to detect XRN1 (magenta) and EDC4 (yellow) by immunofluorescence microscopy (n=2 independent experiments). H. U-2 OS cells were treated as in (E) and immunofluorescence microscopy done to detect DCP1A with the percent cells with PBs from n=3 independent experiments shown (≥318 cells counted per treatment across all replicates). Representative images shown with the average +/- s.e.m. are shown for each experiment with green, gray, and pink points representing the average of each replicate. Statistical significance was assessed with an ordinary one-way ANOVA followed by Tukey’s multiple comparisons test with * p≤ 0.05, ** p≤ 0.01, ***p≤ 0.005, **** p≤ 0.001.

### tRNA synthetase activity is required for stress granule and P-body assembly

While inhibition of tRNA charging activates the ISR, the results of past studies suggest that translation elongation is also inhibited (*62*, *64*). We next tested if suppressing tRNA synthetase activity induced stress granule formation, as halofuginone robustly increased the levels of P-eIF2α. However, stress granule assembly is driven by the release of mRNAs from translation, which would require either ribosomes to run off of mRNAs or ribosome release through RQC/RQT pathways (*11–13*, *48*, *49*, *52*, *58*, *86*). We treated U-2 OS cells expressing the SG marker protein GFP-G3BP1 to mark SGs (*87*) with halofuginone for 1 hour and assessed SG formation. A key observation is that cells treated with halofuginone did not form SGs (**Fig. 1C**). In contrast, cells formed abundant SGs during treatment with canonical stressors arsenite and thapsigargin that led to similar P-eIF2α levels as seen in HF-treated cells (**Fig. 1C**). Furthermore, no SGs were detected in cells treated with halofuginone when visualizing other canonical SG markers including UBAP2L, PABPC1, and polyadenylated RNA (**Fig. 1D**). These results demonstrate tRNA synthetase inhibitors do not cause SG formation despite activating the ISR.

To test whether inhibition of tRNA charging with halofuginone prevents SG assembly under conditions that cause SG formation, we co-treated cells with halofuginone and arsenite or thapsigargin. We observed halofuginone co-treatment prevented SG formation by either arsenite or thapsigargin (**Fig. 1E**). Halofuginone co-treatment did not decrease P-eIF2α induction in response to arsenite (**Fig S1A**) or thapsigargin (**Fig S1B**), indicating inhibition of SGs by halofuginone is not through reduced activation of the ISR. Thus, tRNA synthetase activity is required for SGs to form.

We next determined whether tRNA synthetase inhibition broadly impacts stress-induced cytoplasmic RNP granules by assessing the formation of processing bodies (P-bodies or PBs). P-bodies are enriched in RNA decay factors (e.g., DCP1A, EDC4, XRN1), harbor non-translating mRNAs, and interact with SGs during stress (*7*, *12*, *50*, *88*, *89*). We used immunofluorescence microscopy to detect the PB marker DCP1A in cells treated with halofuginone, arsenite, or thapsigargin. We found that halofuginone inhibited PB assembly (**Fig. 1F-G**). While 22.4+/-2.0% of cells harbor PBs in the unstressed condition, 6.2+/-1.3% of halofuginone-treated cells had PBs (**Fig. 1F**). In contrast, arsenite and thapsigargin induced PBs to 100.0 +/- 0.0% and 95.7 +/- 0.4%, respectively (**Fig. 1F**). Similar results were observed when immunofluorescence was used to detect additional canonical PB proteins, XRN1 and EDC4 (**Fig. 1G**). Therefore, tRNA synthetase inhibition does not induce PB assembly despite activating the ISR.

We next determined whether tRNA synthetase inhibition prevents the assembly of PBs in response to stress. We co-treated cells with halofuginone and arsenite or thapsigargin and measured PBs by staining for DCP1A. We observed that halofuginone significantly decreased the percentage of cells with PBs from 97.9 +/- 1.2% to 52.9 +/- 16.9% in arsenite stress and from 96.9 +/- 0.5% to 7.5 +/- 4.6% in thapsigargin stress (**Fig. 1H**). Together, these results suggest that tRNA charging is required for the assembly of stress-induced RNP granules.

### Inhibition of tRNA charging blocks stress-induced mRNA release from polysomes

Inhibition of translation elongation by halofuginone causes ribosomes to accumulate on proline codons, at the 5’ end of coding sequences, and on mRNAs encoding proline-rich proteins (*62*, *64*). Since we found SG and P-body assembly was blocked despite robust levels of P-eIF2α, we hypothesized that halofuginone would cause retention of mRNAs in polysomes, as ribosomes lack the charged prolyl-tRNA to elongate and allow ribosome runoff. To test this hypothesis, we used polysome profiling to measure mRNA association with ribosomes in lysates of cells treated with halofuginone or the canonical stressor thapsigargin. In support of our hypothesis, we found that unstressed cells exhibited robust polysomes with a high polysome:monosome ratio and thapsigargin treatment caused polysomes to collapse and increased the 80S monosome peak (**Fig. 2A**). In contrast, polysomes were largely preserved in cells treated with halofuginone (**Fig. 2A**). Because halofuginone caused translational repression as measured by metabolic labeling (**Fig. 1B**), these results demonstrate halofuginone causes elongating ribosomes to stall on mRNAs.

**Figure 2.**
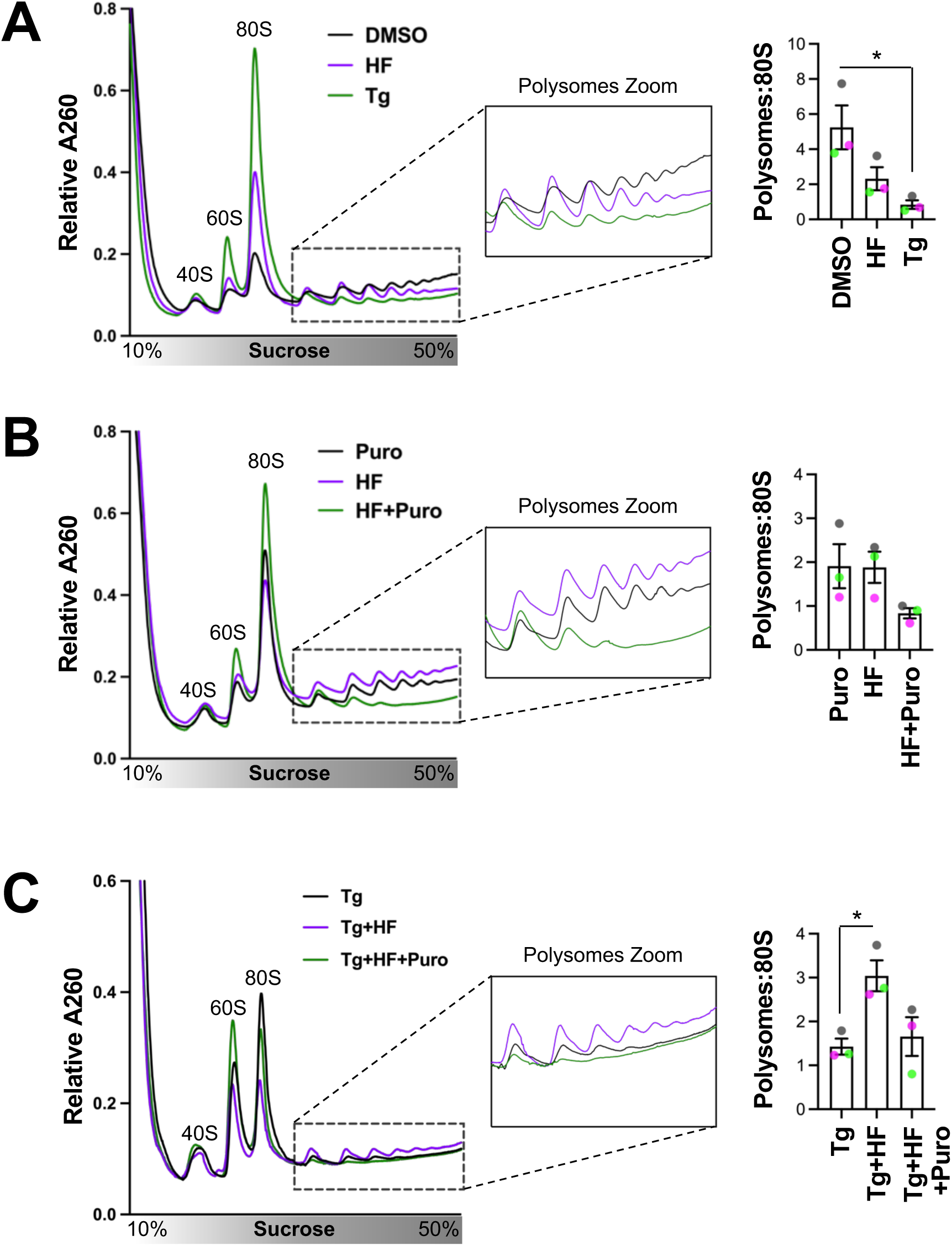
tRNA charging is required for stress-induced mRNA release from polysomes. A. U-2 OS cells were treated with 20 µM halofuginone (HF, purple line), 1.25 µM thapsigargin (Tg, green line), or DMSO carrier (0.2%, black line) for 1 hour and polysome profiling done, with representative profiles at left (n=3 independent experiments). The area under the polysome fractions and 80S peak were quantified and shown at right. B. Cells were treated with 10 µg/mL puromycin (black line), 20 µM HF (purple line), or puromycin (10 µg/mL) and HF (20 µM) (green line) for 1 hour and polysome profiling performed and quantified as in (A) (n=3 independent experiments). C. Cells were treated with Tg (1.25 µM, black line), HF (20 µM) plus Tg (1.25 µM), or HF (20 µM) (purple line) and Tg (1.25 µM) and puromycin (10 µg/mL) (green line) and polysome profiling done and quantified as in (A) (n=3 independent experiments). The average of 3 independent experiments with s.e.m. is shown with the average of each replicate in pink, green, or gray. Significance was assessed with ordinary one-way ANOVAs and Tukey’s multiple comparisons tests with * p<0.05.

We next determined whether puromycin could release mRNAs from stalled ribosomes during halofuginone treatment. Puromycin is a small molecule peptidyl-tRNA mimic that causes premature ribosome release when incorporated into nascent chains (*90*). We anticipated that puromycin would react with empty ribosome A sites awaiting charged Pro-tRNA^Pro^ and cause polysome collapse during halofuginone treatment. We found that puromycin alone did not induce complete polysome collapse after 1 hour (**Fig. 2B**), likely due to continued translation initiation. However, co-treatment of cells with halofuginone and puromycin reduced the polysome fraction, increased the 80S monosome fraction, and decreased the polysome to monosome ratio ∼2.4-fold from 1.9 +/- 0.4 in cells treated with halofuginone to 0.8 +/- 0.1 in cells treated with halofuginone and puromycin (**Fig. 2B**). Further, halofuginone co-treatment stabilized polysomes in cells treated with thapsigargin, yielding a ∼2.1-fold increase in the polysome to monosome ratio from 1.4 +/- 0.2 to 3.0 +/- 0.4 (**Fig. 2C**). These observations suggest that halofuginone causes ribosome stalls to persist in the presence of a canonical stressor. Puromycin reversed the effects of halofuginone on the polysome profile in cells treated with thapsigargin, and restored the polysome to monosome ratio to 1.7 +/- 0.4. These results demonstrate that puromycin rescues ribosome stalling induced by inhibition of tRNA charging by releasing mRNAs from polysomes.

### Ribosome collisions contribute to activation of the ISR by halofuginone

Because polysomes do not collapse and RNP granules do not assemble upon halofuginone stress, we next asked whether ribosome-associated quality control (RQC) pathways failed to resolve stalled ribosomes that result from uncharged tRNA accumulation. We first assessed if halofuginone causes ribosome collisions by measuring ubiquitination of ribosomal protein eS10. The initial stage of the RQC is recognition of the ribosome collision interface and the ubiquitination of ribosomal proteins, including eS10, by the E3 ubiquitin ligase ZNF598 (*91–95*). We observed that treatment with a low concentration of emetine, an irreversible inhibitor of ribosome translocation (*96*) that causes ribosome collisions (*44*, *97*), induced robust ubiquitination of eS10, with a 6.3 +/- 0.5 fold increase over the DMSO carrier control (**Fig. 3A**). A key observation is that halofuginone increased eS10 ubiquitination by 3.1 +/- 0.8-fold relative to DMSO controls at 0.5 hr, the earliest time point collected (**Fig. 3A**). However, Ub-eS10 levels in halofuginone-treated cells did not reach those observed with emetine treatment (**Fig. 3A**), suggesting ribosome collisions occur infrequently in this context. These results suggest that the inhibition of tRNA charging leads to ribosome stalls and a low level of collisions that are recognized and cleared by the RQC pathway.

**Figure 3.**
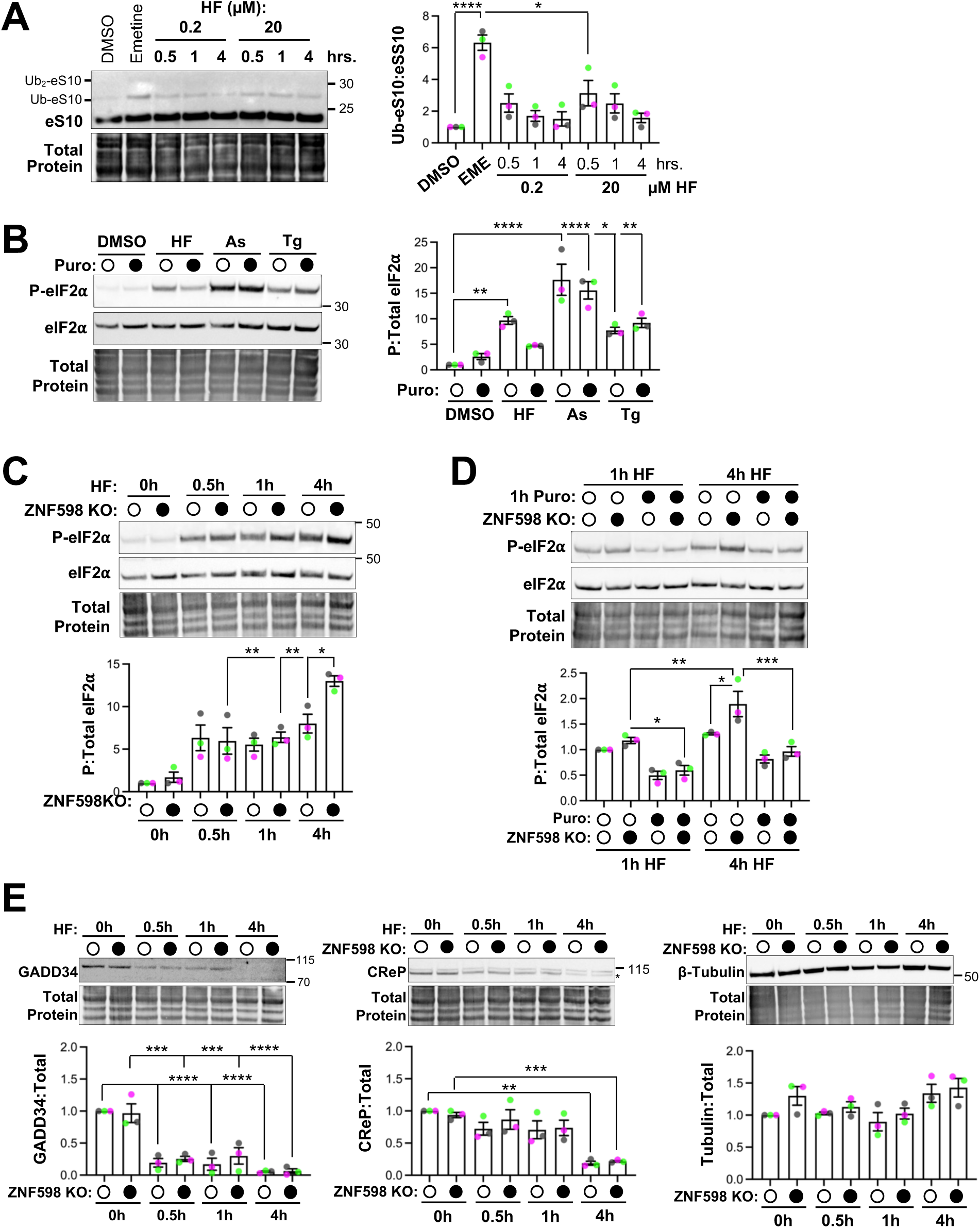
Low-level ribosome collisions contribute to ISR activation during tRNA synthetase inhibition. A. U-2 OS cells were treated with DMSO carrier (0.2%), emetine (0.18 µM) for 15 minutes, or halofuginone (HF, 0.2 or 20 µM) for 0.5, 1, or 4 hours and western blotting for eS10 was done. The higher molecular weight mono-ubiquitylated (Ub-eS10) and unmodified eS10 bands were quantified with average Ub-eS10:eS10 ratios from n=3 independent experiments shown at right and a representative blot with total protein shown at left. B. determined by western blotting. Quantification of P-eIF2α:eIF2α is shown at right, with representative blot with total protein shown at left from n=3 independent experiments. C. Western blotting to total and phosphorylated eIF2α was done from wild-type (WT) and ZNF598 knockout (KO) cells treated with HF (20 µM) for 0.5, 1, or 4 hours. Representative blot with total protein shown (top) and results from n=3 independent replicates quantified (bottom). D. Western blotting to P-eIF2α and total eIF2α from WT and ZNF598 KO cells treated with HF (20 µM) for 1 or 4 hours in the presence or absence of puromycin (10 µg/mL) added 1 hour prior to collection. Representative blot with total protein (top) and quantification of P-eIF2α:eIF2α (bottom) shown from n=3 independent experiments. E. Wild-type and ZNF598 KO cells were treated as in (C) and western blotting for GADD34, CReP (marked by an asterisk), and β-tubulin was done with quantification done relative to total protein. Representative blots shown at top and at bottom are results from n=3 independent experiments. Molecular weights (kDa) are indicated on each blot. Quantification is reported as average +/- s.e.m. with pink, gray, and green points representing each replicate. Statistical significance was determined with ordinary one-way ANOVAs and Tukey’s multiple comparisons tests, with * p≤ 0.05, ** p≤ 0.01, *** p≤ 0.005, **** p≤ 0.001.

We next asked whether ribosome collisions contribute to activation of the ISR by halofuginone. Uncharged tRNAs can activate the ISR via GCN2 (*38*, *39*, *41*, *42*, *64*, *98*, *99*). We assessed P-eIF2α levels in cells treated with halofuginone, arsenite, or thapsigargin in the presence or absence of puromycin to prevent ribosome collisions by releasing mRNAs from ribosomes (*38*). We found that while halofuginone significantly increased P-eIF2α levels, the addition of puromycin reduced the levels of P-eIF2α by ∼2-fold in cells co-treated with halofuginone (**Fig. 3B**). There were no significant differences in P-eIF2α levels upon puromycin co-treatment with arsenite or thapsigargin. These results suggest that ribosome occupancy contributes to, but is not solely responsible for, activation of the ISR after loss of tRNA charging.

Since the RQC pathway resolves ribosome collisions, we hypothesized that activation of RQC by halofuginone would limit GCN2 activation and reduce P-eIF2α levels. We reasoned that P-eIF2α levels resulting from collisions would be elevated in cells that cannot clear collided ribosomes due to genetic depletion of ZNF598, which initiates the RQC pathway (*94*). We generated a genetically depleted ZNF598 knockout U-2 OS cell pool (ZNF598 KO) using CRISPR/Cas9. We verified ZNF598 was knocked out by performing Sangar sequencing of the genomic DNA (**Fig. S2A**) and ICE analysis (**Fig. S2B**). Further, while western blotting with a ZNF598 antibody showed a residual expression of a lower molecular weight protein in the knockout pool, emetine treatment did not cause Ub-eS10 levels to increase in these cells, suggesting this lower molecular weight ZNF598 protein may be a non-functional in-frame deletion (**Fig. S2C**). We then assessed P-eIF2α levels in wild-type and ZNF598 KO cells treated with halofuginone at 0.5, 1, and 4 hr. At 4 hr post-halofuginone treatment, ZNF598 KO cells had a significant, 1.63-fold increase in P-eIF2α compared to wild-type cells (**Fig. 3C**). No significant differences in P-eIF2α were observed early in stress (**Fig. 3C**). Therefore, ZNF598 limits activation of the ISR when tRNA charging is inhibited.

We hypothesized that the observed increase in P-eIF2α levels during halofuginone treatment in the ZNF598 KO cell line was due to the accumulation of ribosome collisions, which would otherwise be cleared by the RQC pathway. We performed two experiments to test this hypothesis. First, we assessed whether stalled ribosomes contributed to the elevated P-eIF2α levels observed in ZNF598 KO cells. We co-treated wild-type and ZNF598 KO cells with puromycin during 1 or 4 hr of halofuginone stress to resolve stalled ribosomes and assessed P-eIF2α levels. We found that puromycin reduced P-eIF2α levels in ZNF598 KO cells to levels comparable to those observed in wild-type cells treated with halofuginone alone (**Fig. 3D**). This result suggests that the increased P-eIF2α levels in ZNF598 KO cells resulted from persistent collided ribosomes that would otherwise be cleared by the RQC pathway.

Second, we assessed how the levels of CReP and GADD34, which interact with protein phosphatase 1 to dephosphorylate P-eIF2α, contribute to P-eIF2α levels throughout the duration of HF treatment. We reasoned that ZNF598 KO cells could display an increase in P-eIF2α during late (4 hr) halofuginone stress compared to wild-type cells if CReP and GADD34 were depleted more in the absence of ZNK598 at that time point. Wild-type cells would otherwise have sufficient P-eIF2α phosphatase activity to limit P-eIF2α levels during halofuginone stress. Our data rules this possibility out, as we found that GADD34 and CReP are equally reduced during halofuginone stress in ZNF598 KO and wild-type cells (**Fig. 3E**). The observed reduction in CReP and GADD34 levels is in line with the results of prior studies that demonstrated CReP is rapidly depleted upon translation suppression (*38*) and GADD34 and CReP are not expressed in cells treated with high levels of halofuginone (*64*). Not all proteins were depleted at 4 hours post-halofuginone treatment, as tubulin and total protein levels were unchanged over time (**Fig. 3E**). Thus, CReP and GADD34 levels are equal in both cell lines throughout the duration of treatment regardless of the presence of ZNF598. Taken together, these results demonstrate that ribosome collisions occur during tRNA synthetase inhibition, cause an increase in P-eIF2α levels, and do so via increased phosphorylation of eIF2α and not through decreased phosphatase levels.

### Inducing ribosome release rescues RNP granule assembly after tRNA synthetase inhibition

We next asked if unresolved ribosome stalls block RNP granule assembly when tRNA synthetase activity is inhibited during stress. First, we assessed whether stabilizing ribosomes on mRNAs during translation elongation prevented stress-induced RNP granule assembly. Consistent with past research (*11–13*, *48*, *49*), trapping mRNAs in polysomes with the irreversible ribosome translocation inhibitor emetine (*96*), prevents SGs from forming during arsenite stress (**Fig. 4A**). Emetine also significantly reduced PBs in unstressed cells, but did not impact their assembly during arsenite stress, similar to prior results of experiments with the translation elongation inhibitor cycloheximide (*11*, *12*, *50*, *89*, *100*) (**Fig. 4A**). Therefore, blocking translation elongation inhibits stress granule and constitutive P-body assembly.

**Figure 4.**
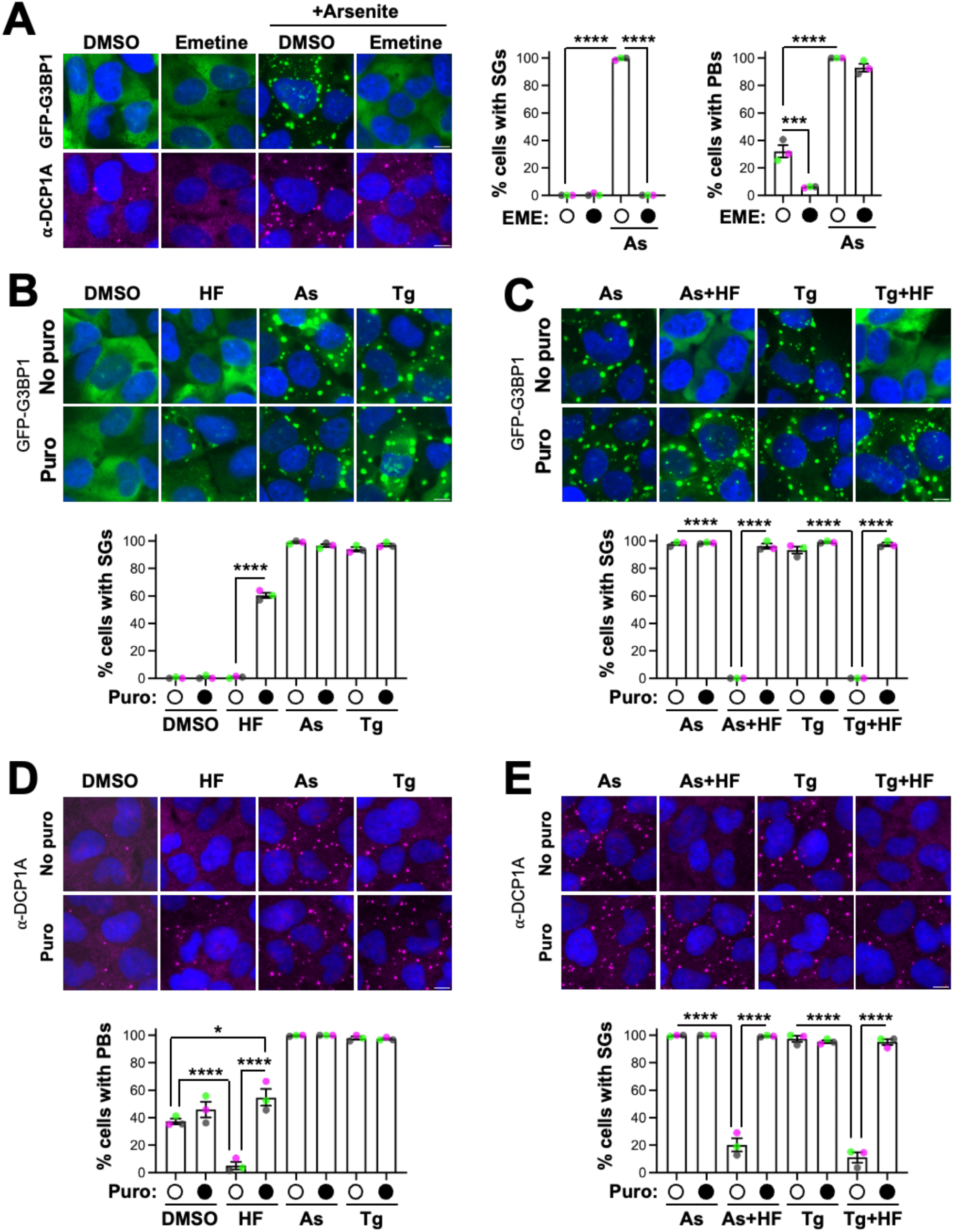
Ribosome release rescues stress-induced RNP granule assembly upon tRNA synthetase inhibition. A. U-2 OS cells stably expressing GFP-G3BP1 were treated with emetine (180 µM) or DMSO (0.2%) with sodium arsenite (As, 250 µM) for 1 hour. Immunofluorescence microscopy was performed for P-body (PB) marker DCP1A and the SG marker GFP-G3BP1 was imaged. The percent cells with SGs or PBs was quantified, with ≥349 cells counted per treatment across all replicates (shown at right) from n=3 independent experiments. B. Cells were treated with DMSO (0.2%), 20 µM HF, 250 µM As, or 1.25 µM thapsigargin (Tg) with or without puromycin (10 µg/mL) for 1 hour, followed by imaging and quantification of SGs. The percent cells with SGs from n=3 independent experiments is shown (bottom) with representative images above. ≥388 cells counted per treatment across all replicates. C. Cells were treated with HF (20 µM) in replicates. D. Cells were treated as in (B) and IF for DCP1A performed. The percent cells with P-bodies (PBs) from n=3 independent experiments is shown at bottom with representative images at top, with ≥342 cells counted per treatment across all replicates. E. Cells were treated as in (C) and IF performed to detect DCP1A. The percentage of cells with PBs was quantified from n=3 independent experiments (at bottom) with representative images shown (at top). ≥378 cells counted per treatment across all replicates. Quantification is reported as average +/- s.e.m., with green, pink, or gray dots indicating the average of each replicate. All scale bars are 10 µm. Statistical significance was assessed with an ordinary one-way ANOVA followed by Tukey’s multiple comparisons test: * p ≤ 0.05, *** p≤ 0.005, **** p≤ 0.001.

To directly test whether ribosome stalling during tRNA synthetase inhibition prevents RNP granule assembly, we used puromycin to release mRNAs from ribosomes during halofuginone treatment and evaluated SG and PB assembly. While SGs were not observed in halofuginone treatment alone, co-treatment with puromycin and halofuginone for 1 hour caused SG assembly in 60.5 +/ 2.0% of cells (**Fig. 4B**). Puromycin did not cause SG assembly in unstressed cells or alter SG assembly upon treatment with arsenite or thapsigargin (**Fig. 4B**). Similarly, while SGs did not form in the presence of halofuginone during either arsenite or thapsigargin stress, puromycin co-treatment rescued SG assembly (**Fig. 4C**).

We next determined if P-bodies are also limited by mRNA retention in polysomes upon halofuginone treatment. Puromycin co-treatment significantly increased the percentage of cells with PBs, from 5.1 +/- 2.7% in cells treated with halofuginone alone to 54.8 +/- 6.1% in co-treated cells (**Fig. 4D**). In contrast to what was observed with emetine treatment, halofuginone blocked the assembly of P-bodies upon arsenite or thapsigargin stress (**Fig. 4E**). Co-treatment with puromycin also rescued PB assembly in cells treated with halofuginone and arsenite or thapsigargin (**Fig. 4E**). Specifically, the percentage of cells with PBs increased from 20.2 +/- 4.8% in cells with arsenite and halofuginone to 99.4 +/- 0.4% when puromycin was added, and from 11.1 +/- 3.8% in cells with thapsigargin and halofuginone to 95.1 +/- 2.2% upon addition of puromycin. Together, these results suggest that RNP granules do not assemble when tRNA charging is impaired in part because stalled ribosomes remain associated with mRNAs.

One alternative possibility is that puromycin drives RNP granule assembly by increasing ISR activation in halofuginone-treated cells. Puromycin can induce SGs at high concentrations or long treatment duration (i.e. 24 hours) (*13*, *101*, *102*) and increases P-eIF2α levels in association with reduced CReP expression (*38*). We found Ub-eS10 levels suggestive of ribosome collisions are elevated early in halofuginone stress (**Fig. 3A**) at similar times when GADD34 and CReP are at their highest levels during halofuginone treatment (**Fig. 3E**). If ribosome collisions contribute to P-eIF2α levels early during halofuginone treatment, we reasoned that puromycin co-treatment would reduce P-eIF2α levels by resolving stalled, collided ribosomes that activate GCN2. We therefore assessed P-eIF2α levels in cells treated with halofuginone in the presence or absence of puromycin over a 4 hour time course. Consistent with the idea that puromycin releases stalled, collided ribosomes from mRNAs that contribute to ISR activation, we observed puromycin reduced P-eIF2α levels early, but not late, during halofuginone treatment (**Fig S3A**). We also assessed stress granule assembly over the same time course of halofuginone treatment in the presence or absence of puromycin. A key observation is that fewer cells formed SGs early during halofuginone treatment versus longer-term treatment (**Fig. S3B**). The percent of cells with SGs increased over time in cells treated with halofuginone and puromycin, from 35.5 +/- 2.0% of cells with SGs after 1 hour of co-treatment to 94.7 +/- 2.5% of cells after 4 hours of co-treatment (**Fig. S3B**). Together, these observations are consistent with the idea that puromycin minimizes ISR activation early in halofuginone stress and limits SG assembly by preventing ribosome collisions, while promoting SG assembly later during halofuginone stress by releasing mRNAs from stalled ribosomes.

Halofuginone specifically inhibits the prolyl-tRNA synthetase activity of EPRS1 (*36*) and does not affect the extent of charging of any of the other 19 tRNA isoacceptors (*38*). We therefore tested whether inhibiting tRNA charging of other amino acids also blocked the assembly of SGs and PBs through ribosome stalling. To do so, we treated cells with borrelidin, which inhibits threonyl-tRNA synthetase (*103*) and also activates the ISR via GCN2 (*104*). We found that borrelidin induced similar levels of P-eIF2α as thapsigargin, with a 10.4 +/- 1.4 fold increase of borrelidin over control versus a 11.1 +/- 0.6 fold increase with thapsigargin stress (**Fig. S4A**). Consistent with our previous findings with halofuginone, no SGs were observed in cells treated with borrelidin, and borrelidin inhibited thapsigargin-induced SGs (**Fig. S4B**). Further, puromycin co-treatment induced SG assembly in cells treated with borrelidin (to 77.9 +/- 4.6% of cells), and rescued SG assembly in cells stressed with thapsigargin and borrelidin (**Fig. S4C**). Similarly, the percentage of cells with PBs was decreased from 30.4 +/- 7.4% in unstressed cells to 2.8 +/- 0.8% during borrelidin treatment, and was restored to 20.9 +/- 4.9% by puromycin (**Fig. S4D**). Borrelidin also decreased the percentage of cells with PBs during thapsigargin treatment from 99.3 +/- 0.7% to 5.1 +/- 1.9%, an effect that was rescued to 94.8 +/- 1.9% by puromycin. Therefore, the uncoupling of RNP granule assembly from the ISR is generalizable across multiple tRNA charging inhibitors.

### tRNA synthetase inhibition causes persistent ribosome stalls

Multiple cellular mechanisms exist to resolve ribosomes stalled in elongation, including ribosome-associated quality control (RQC) and ribosome rescue factors related to the translation termination machinery (*105*, *106*). We therefore hypothesized that ribosome stalls caused by tRNA charging inhibition would be removed from mRNAs, enabling RNP granule assembly. We tested this hypothesis by determining if SGs formed during long-term tRNA charging inhibition using live cell imaging of the SG marker GFP-G3BP1 in cells treated with halofuginone. We found that halofuginone-treated cells did not form SGs for up to 16 hours of treatment, the limit of imaging before cytotoxicity was observed (**Fig. 5A**). The absence of SGs was not due to detoxification of halofuginone or resolution of stress, as P-eIF2α levels remained high up to 16 hours (**Fig. 5B**). Additionally, polysome profiling confirmed that polysomes persist at 16 hours of stress (**Fig. 5C**). These results suggest that ribosome stalls due to the loss of tRNA charging are cleared either very slowly or not at all.

**Figure 5.**
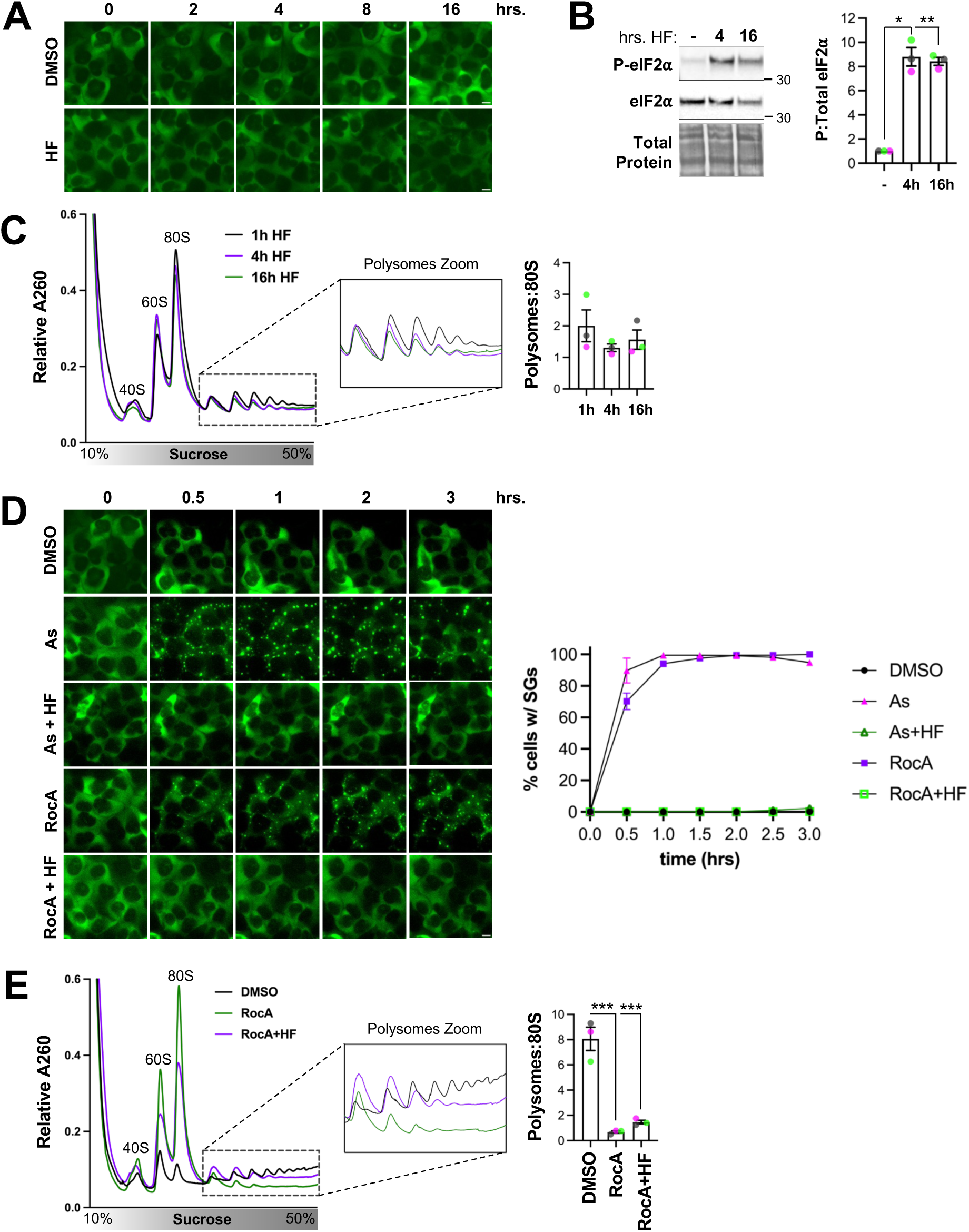
tRNA synthetase inhibition causes persistent ribosome stalls. A. U-2 OS cells expressing GFP-G3BP1 were treated with halofuginone (HF, 20 µM) or DMSO carrier (0.2%) and images collected every 2 hours for 16 hours, representative images shown (n=3 independent experiments). B. Cells were treated with DMSO (-) or HF (20 µM) for 4 or 16 hours, and P-eIFα and total eIF2α detected by western blotting. Representative blots with total protein shown at left with average +/- s.e.m. of n=3 independent experiments shown at right. Molecular weights (kDa) shown on each blot. C. Cells were treated with HF (20 µM) for 1 (black line), 4 (purple line), or 16 (green line) hours and polysome profiles performed. Representative profile shown at left with the average +/- s.e.m. of the polysome:monosome ratio from n=3 independent replicates shown at rocaglamide A (RocA, 2 µM), and GFP-G3BP1 imaged every 0.5 hours for 3 hours. Representative images shown at left with the average +/- s.e.m. percent cells with SGs from n=3 independent experiments shown at right (≥301 cells counted per treatment across all replicates). E. Cells were treated with DMSO (0.2%) (black line), RocA (2 µM) (green line), or RocA (2 µM) plus HF (20 µM) (purple line) for 3 hours followed by polysome profiling with representative profiles shown at left and the average +/- s.e.m from n=3 independent experiments shown at right, with pink, gray, and green dots representing each replicate. All scale bars are 10 µm. Significance was assessed with ordinary one-way ANOVAs and Tukey’s multiple comparisons tests with * p≤ 0.05, ** p≤ 0.01, *** ≤ 0.005.

One alternative possibility is that ribosome stalls are cleared, but are also slowly replaced by residual translation initiation activity. To test this possibility, we performed live cell imaging of GFP-G3BP1-expressing cells treated with halofuginone and either arsenite or rocaglamide A, which inhibit translation initiation via the ISR or eIF4A inhibition (*107*), respectively. We found that while robust SG assembly occurred in rocaglamide A- or arsenite- treated cells, halofuginone completely prevented SG formation in cells co-treated with rocaglamide A or arsenite up to 3 hours later, the duration of imaging (**Fig. 5D**). In line with these results, halofuginone limited polysome collapse upon rocaglamide A treatment (**Fig. 5E**). Specifically, after 3 hours of treatment the polysome to 80S ratio was increased ∼2.1-fold, from 0.7 +/- 0.1 in cells treated with rocaglamide A alone to 1.5 +/- 0.2 in co-treated cells with rocaglamide A and halofuginone. These results suggest that the ribosomes associated with mRNAs after long-term halofuginone treatment are stable, stalled species, rather than the result of continued translation initiation.

## Discussion

Here we show that tRNA charging is essential for stress-induced RNP granule assembly by preventing persistent, unresolved ribosome stalls. First, we show tRNA synthetase inhibitors induce the ISR and suppress translation without driving the assembly of stress-induced RNP granules. We demonstrate that prolyl- and threonyl- tRNA synthetase inhibitors block the assembly of SGs and PBs during stress by trapping mRNAs in stalled polysomes. We observed low-level activation of the ribosome-associated quality control pathway initiated by ZNF598, suggesting ribosome collisions occur and are cleared early upon tRNA synthetase inhibition. The results of experiments monitoring P-eIF2α levels in wild-type and ZNF598-depleted cells in the presence or absence of puromycin suggest ribosome collisions contribute to ISR activation when tRNA charging is inhibited. Residual stalled ribosomes that accumulate during tRNA synthetase inhibition persist and prevent RNP granule assembly, suggesting they are not rescued by RQC pathways. Taken together, we find RNP granules are uncoupled from the stress response when tRNA charging is inhibited due to persistent ribosome stalls (**Fig. 6**).

**Figure 6.**
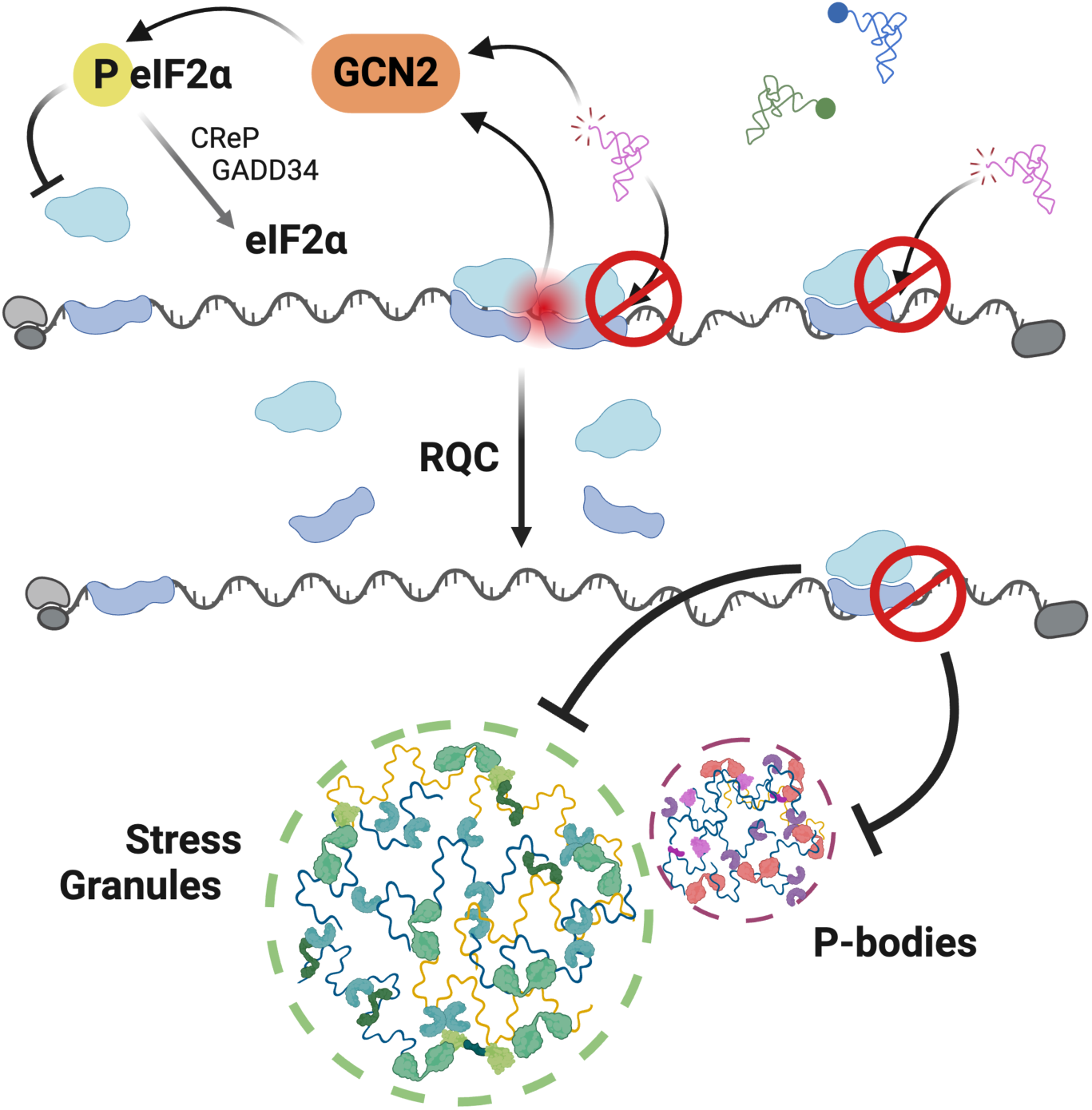
Model depicting the requirement of tRNA synthetase activity for RNP granule assembly. Upon tRNA synthetase inhibition, ribosomes stall and eIF2α phosphorylation results from uncharged tRNAs and collided ribosomes. Ribosome-associated quality control (RQC) clears stalled, collided ribosomes but cannot resolve ribosomes that stall in the absence of collisions. The association of stalled ribosomes with mRNAs blocks the assembly of stress granules and P-bodies.

One implication of these results is that other stressors that activate the ISR by disrupting translation elongation may also inhibit RNP granule assembly because of persistent ribosome stalls. Chemical agents that damage RNA such as methyl methanesulfonate (MMS) (*32*, *34*), nitric oxide (*35*), and 4-nitroquinoline 1-oxide (*32*) cause ribosome stalling and ISR activation. MMS has been reported to not induce SGs (*108*, *109*), and nitric oxide triggers the formation of atypical, stable SGs that assemble more rapidly in the presence of puromycin (*110*). Further, chemotherapeutic agents such as etoposide and cisplatin induce tRNA degradation via SLFN11 nuclease, causing ribosome stalling and GCN2 activation (*31*). Etoposide has been reported to not cause SG assembly (*109*), while cisplatin causes aberrant, small SGs to form in only ∼20% of cells despite robust translation suppression, and inhibits the formation of arsenite-induced SGs (*111*, *112*). Ribosome-stalling stressors may take divergent routes to SG assembly when polysome-free mRNA is unavailable to seed or scaffold them. While cisplatin-induced stress granule-like assemblies were reported to not be stimulated by puromycin treatment (*111*), cisplatin-induced “stress granules” have no (*111*) or low (*111*, *112*) levels of poly(A) mRNA (*111*) and persist after stress removal (*111*, *112*), suggesting they are not canonical SGs. Similarly, UV stress causes mRNA damage, ribosome stalling, and ISR activation via GCN2 (*32*, *44*, *113*) but induces few SGs to form (*114*). These UV-induced SGs are not enriched in PABP or translation factors found in canonical SGs (e.g., eIF4G, eIF3B, 4E-BP, S6) (*114*) or poly(A) mRNA (*115*), and may instead be scaffolded by damaged RNAs released from the nucleus (*116*). One possibility is that this unusual mechanism of SG assembly occurs because polysome-free mRNAs are unavailable. Indeed, a recent study showed that UV stress causes ∼2.3-fold slower ribosome runoff compared to untreated controls and that nuclease-resistant collided ribosomes persist for at least 4 hours (*32*). Therefore, stress-induced ribosome stalling that limits RNP granule assembly may occur in diverse stress conditions.

Our results predict that a key determinant of whether elongation-targeting stressors will induce or prevent RNP granules is if the ribosome stalls they generate can be recognized and cleared by ribosome release factors and/or RQC. Here, we observed low-level activation of the RQC pathway upon robust inhibition of tRNA charging. The observed ubiquitination of ribosomal protein eS10 by ZNF598 upon halofuginone treatment is suggestive of ribosome collisions, as this initiator of the RQC uniquely recognizes the disome interface (*54*, *92*). We therefore anticipated that stalled ribosomes would be rapidly cleared from mRNAs by the RQC/ribosome quality trigger (RQT) complex downstream of ZNF598 activity (*54*, *92*, *117*). The results of single molecule live cell imaging assays of reporter mRNAs harboring a poly(A)-tract in the coding region that cause ribosome collisions demonstrated that ∼50% of the mRNAs are released from stalled ribosomes within 15 minutes (*95*). However, we observed that halofuginone-induced ribosome stalls persisted for over 16 hours. One possible explanation is that ribosome collisions make up a minor species among the ribosome stalls generated by severe inhibition of tRNA charging. For instance, proline accounts for 6.3% of all amino acids in the human proteome (*118*), and ribosome stalling at every proline codon could cause a similar effect as using high concentrations of elongation inhibitors that freeze ribosomes in isolated stalls rather than in collided disomes (*44*, *119*). Therefore, inhibition of tRNA synthetases may cause isolated ribosome stalls that go unrecognized by RQC/RQT pathways.

We made several observations that suggest the RQC pathway is activated during halofuginone stress and indicate that ribosome collisions contribute to the ISR upon acute tRNA synthetase inhibition. The observed accumulation of ubiquitinated eS10 suggests ribosome collisions occur early during acute, high level halofuginone treatment. Further, our observation that puromycin reduces P-eIF2α levels at early time points during halofuginone stress indicates stalled ribosomes may contribute to ISR activation. We considered the possibility that puromycin reduces P-eIF2α levels by interacting with GCN2 and outcompeting interactions with uncharged tRNAs that accumulate with tRNA synthetase inhibitors. However, GCN2 preferentially binds to uncharged tRNAs (*39*) and puromycin mimics an aminoacylated (tyrosyl) tRNA. Additionally, we found that cells deficient in the ribosome collision sensor ZNF598 have elevated P-eIF2α levels compared to wild-type cells during halofuginone treatment, although this increase in P-eIF2α was only apparent when the P-eIF2α phosphatases are depleted (likely due to halofuginone-induced translational suppression (*64*)).

Our results are in line with recent work demonstrating that the ISR and RQC pathways are antagonistic (*34*, *44*, *113*). Chemical inhibition of the ISR with ISRIB increases RNase-resistant disomes and P-eIF2α levels during amino acid starvation (*44*), and GCN2 depletion increases Ub-eS10 levels and collided ribosomes during UV-C stress (*113*) by increasing translation initiation and ribosome load during elongation stresses. In yeast, loss of Hel2 (the yeast ortholog of ZNF598) or the ribosome release factor Slh1 (the yeast ortholog of ASCC3) increases Gcn2 activation, and Gcn2 depletion increases MMS-induced protein ubiquitination that likely represent ubiquitinated ribosomal proteins (*34*). Finally, recent work suggests Gcn2 is preferentially activated by stalled ribosomes with empty A sites (*34*), which likely accumulate upon tRNA synthetase inhibition. Ribosome profiling analysis demonstrates halofuginone causes ribosomes to stall with proline codons at the ribosomal A site, suggesting ribosomes are awaiting an incoming prolyl-tRNA (*62*). Dimethyl sulfate probing and ribosome profiling experiments revealed that depletion of glutamyl-tRNA, histidine biosynthesis inhibitors, and glutamine starvation all result in accumulation of 21 nt ribosome footprints indicative of an empty A-site (*120*). While a recent study (*38*) demonstrated that puromycin does not reduce P-eIF2α levels in wild-type HEK293T cells treated with nanomolar levels of halofuginone, this analysis was performed after a 3 hour treatment when collided ribosomes may have already been cleared. Therefore, our results suggest that robust tRNA synthetase inhibition partially activates the ISR through ribosome collisions.

Uncoupling of RNP granule assembly from the ISR by elongation stressors may affect post-transcriptional gene regulation, cell signaling, and cellular resilience. It is estimated that ∼5 - 15% of cytoplasmic polyadenylated RNA localizes to SGs (*10*, *14*, *121*, *122*), and non-translating mRNAs can remain associated with SGs and PBs for long durations suggesting they are sequestered within them (*7–9*, *52*). Furthermore, genome-wide analyses demonstrate differential enrichment of mRNAs within SGs and PBs, with preferential localization of long RNAs and mRNAs that are translationally repressed to these RNP granules (*14–21*). While it is not clear whether localization to a SG or PB impacts the translation or degradation of mRNAs after stress (*88*, *123–128*), sequestration of non-translating mRNAs within these RNP granules limits their access to the translating pool and may change their structure and/or interaction partners to impact gene expression during and after stress. The results of several studies suggest SGs can act as hubs of signaling pathways that promote cell survival by harboring signaling proteins (reviewed in (*129*)). The sequestration of the signaling scaffold protein RACK1 within SGs is associated with suppression of apoptosis (*109*). Additionally, the mTORC1 protein Raptor localizes to SGs in association with reduced mTORC1 activity (*130*), and DYRK3 kinase is thought to mediate mTORC1 activation by promoting SG disassembly (*131*). However, stress granules and P-bodies have been implicated as beneficial (*22–26*) or detrimental (*27–30*), in diverse cellular and organismal contexts (*132*). Therefore, one implication of our work is that the lack of RNP granule assembly upon tRNA synthetase inhibition could impair gene expression and/or cell signaling events during stress and impact cellular resilience.

tRNA synthetase inhibitors have been extensively investigated as therapies for a wide range of conditions including fibrosis, autoimmunity, cancer, inflammatory diseases, and fungal, bacterial, mycobacterial, and protozoal infections (*133–135*). Borrelidin has anti-malarial properties, and borrelidin analogs show promise in selectively inhibiting *Plasmodium* threonyl-tRNA synthetase activity in a mouse model of malaria and *in vitro* (*136*). Halofuginone and other prolyl-tRNA synthetase inhibitors exhibit anti-fibrotic effects and have been investigated for treatment of systemic sclerosis, graft-versus-host disease, and cardiac fibrosis (*68*, *137*). Halofuginone was designated as an orphan drug for scleroderma and Duchenne muscular dystrophy. While the translation of proline-rich extracellular matrix proteins that are critical for fibrosis such as collagen is likely directly suppressed by halofuginone (*36*, *70*), halofuginone also inhibits the differentiation of T cell subtypes (T_H_17 cells) that can contribute to fibrosis and autoimmunity (*37*, *138*). Halofuginone and borrelidin also exhibit anti-cancer (*64*, *69*, *104*, *139*), anti-inflammatory (*37*, *140*, *141*), and anti-diabetic (*71*, *142*) properties. Future work examining whether and how the activation of an ISR without RNP granule assembly contributes to the therapeutic effects of tRNA synthetase inhibitors may suggest pathogenic mechanisms and new treatment approaches.

Chemical inhibition of tRNA charging may suggest mechanistic insight into genetic diseases caused by loss-of-function mutations in tRNA synthetase genes and genes important for tRNA metabolism. Mutations in six tRNA synthetases are associated with the rare dominant peripheral neuropathy Charcot-Marie-Tooth disease (*72–77*), and mutations in all 19 cytoplasmic tRNA synthetase genes are associated with multisystem disorders (*78*, *79*). Animal models expressing tRNA synthetase mutations associated with disease exhibit increased P-eIF2α levels and/or stress-induced gene expression (*83*, *84*, *143*, *144*), and genetic ablation of the ISR via GCN2 knockout mitigates disease-associated phenotypes (*84*). However, GCN2 knockout increased disease-associated neurodegenerative phenotypes in mice that lack the tRNA^ARG^ isodecoder *n-Tr20* and *Gtpbp1* or *Gtpbp2* (*45*, *57*). Further, cultured fibroblasts from patients harboring compound heterozygous mutations in the prolyl-tRNA charging domain of EPRS1 do not have a constitutive ISR, but rather hyper-activate the ISR in response to ER stress and have reduced viability during chronic ER stress (*80*). Additionally, variants in genes that encode tRNA modification or maturation enzymes (*145*, *146*) may activate the stress response and reduce translation activity (*147*). Cell culture and animal models of genetic diseases with mutations in, or depletion of, tRNA modification enzymes (*148*, *149*), splicing factors (*150*), and RNA polymerase III (*151*) exhibit markers of the stress response. Future work investigating the impact of disease-associated tRNA, tRNA metabolism, and tRNA synthetase gene variants on the ISR, ribosome stalling, and RNP granules may lend insight into disease mechanisms and inform therapeutic strategies.

A broader implication of our work is that persistent stalled, but not collided, ribosomes are tolerated without activating the stress response or being resolved. These results are a departure from the current model that mRNAs trapped in stalled ribosomes are sensed and released by translation quality control machinery. Ribosome stalling without ISR activation or ribosome release may be important in diverse biological contexts. First, local translation is critical for neuronal function, as mRNAs are transported to distal cellular processes within transport granules while they are stalled in translation elongation (*152–158*). In oligodendrocytes, mRNAs are also transported in a translationally repressed state, and the presence of translation factors within these transport granules suggests the possibility that these mRNAs are also stalled in elongation (*159–161*). Second, paused or slowed translation elongation when the ribosome encounters specific mRNA or nascent protein features is critical for co-translational protein folding, translation fidelity, and nascent protein quality control to limit production of misfolded or aggregation-prone proteins (*162*, *163*). Translation elongation rates are largely governed by the levels of cognate tRNAs, amino acid properties, and competition for cognate decoding by tRNAs capable of wobble decoding (*164–166*), suggesting that ribosome stalling as a result of charged tRNA deficiency is ignored by RQC pathways to enable proper protein production. Third, cells may have evolved to limit the release of mRNAs from translation via RQC in stress conditions when nutrients and other resources are low, as releasing mRNAs from the translating pool to later re-initiate translation would require more energy and resources than restarting elongation (*167*). In support of this idea, amino acid deprivation causes ribosomes to accumulate at their corresponding codons (*168*). Finally, as many tRNA synthetase inhibitors are natural products, one possibility is that ribosome stalling is toxic in part because animals have not evolved mechanisms to release uncollided ribosomes. Therefore, the cell may benefit from not recognizing or clearing stalled ribosomes that do not collide to enable mRNA transport, co-translational protein folding and quality control, or recovery from stress conditions.

## Materials & Methods

### Cell culture and treatments

U-2 OS cells stably expressing EGFP-G3BP1 via lentivirus transduction were described previously (*87*) and kindly shared by James Burke and Roy Parker. Cells were cultured in high glucose DMEM with glutamine and pyruvate supplemented with 9% FB essence (FBE; Avantor), 2 mM Glutamax (Gibco) and 1% penicillin-streptomycin in a 5% CO_2_ 37°C incubator. Hoechst staining was done periodically to confirm cells were not contaminated with *Mycoplasma*. The following chemical treatments were performed for the indicated times using: DMSO (0.2%), ethanol (1%), halofuginone (MedChem Express; MCE #HY-N1584) in DMSO at 20 µM or 0.2 µM, sodium arsenite (Ricca Chemical Company) at 250 µM, Borrelidin (MCE) in ethanol at 100 µM for 4 hour, thapsigargin (AG Scientific) in DMSO at 1.25 µM, rocaglamide A (MCE) at 2 µM, puromycin (Gold Biosciences) at 10 μg/mL, and emetine (CalBioChem) in DMSO at 0.18 µM or 180 µM.

To generate ZNF598 knockout cells, Cas9-sgRNA RNP particles were prepared according to manufacturer instructions using a Synthego Gene Knockout Kit to the ZNF598 gene (sgRNA targets: CGCCTTCCGCACCGAGATCG, GAACCGCCACATCGACCTGC, GAACGAGGGTGAGCAGGCAC). EGFP-G3BP1 U-2 OS were nucleofected two times (iteratively) using a Lonza 4D-Nucleofector X unit. Deletions were confirmed using PCR to ZNF598 genomic DNA (Primer sequences: AGTGGTACTCGCGCAAGGACCT, TCCCTTCCCACTGCTCCTGTGG) and Sanger sequencing with alignment to the ZNF598 genomic DNA (ENSG00000167962) sequence using Benchling (Benchling [Biology Software]. (2024). Retrieved from https://benchling.com). Knockout efficiency was estimated from these data using the Inference of CRISPR Edits (ICE) tool (Synthego Performance Analysis, ICE Analysis. 2019. v3.0. Synthego, accessed March 2024) with 100% indel detection and 94% knockout score determined for the cell pool.

### Fluorescence Microscopy

To image RNP granules, U-2 OS cells were grown in glass bottom 96 well plates, treated as indicated, and then fixed for 10 minutes with 4% paraformaldehyde in PBS. To image stress granules using GFP-G3BP1, fixed cells were washed in PBS once, incubated in PBS with NucBlue Live Cell Stain (Hoechst 33342; Invitrogen) for 30 minutes at room temperature, rinsed once, and then imaged in PBS. Immunofluorescence microscopy was done to image endogenous stress granule markers and P-body markers. After fixation, cells were simultaneously permeabilized and blocked for 10 minutes at room temperature in AbDil buffer (PBS with 6% BSA and 0.5% Triton X-100). Primary antibody incubations (see **Table 1**) were performed in 0.5X AbDil in PBS for either 1 hour at room temperature or overnight at 4°C. After washing in PBS three times for 5 minutes each, cells were incubated with secondary antibody in 0.5X AbDil with NucBlue for 1 hour at room temperature, washed three times for 10 minutes each, and imaged in PBS.

**Table 1:**
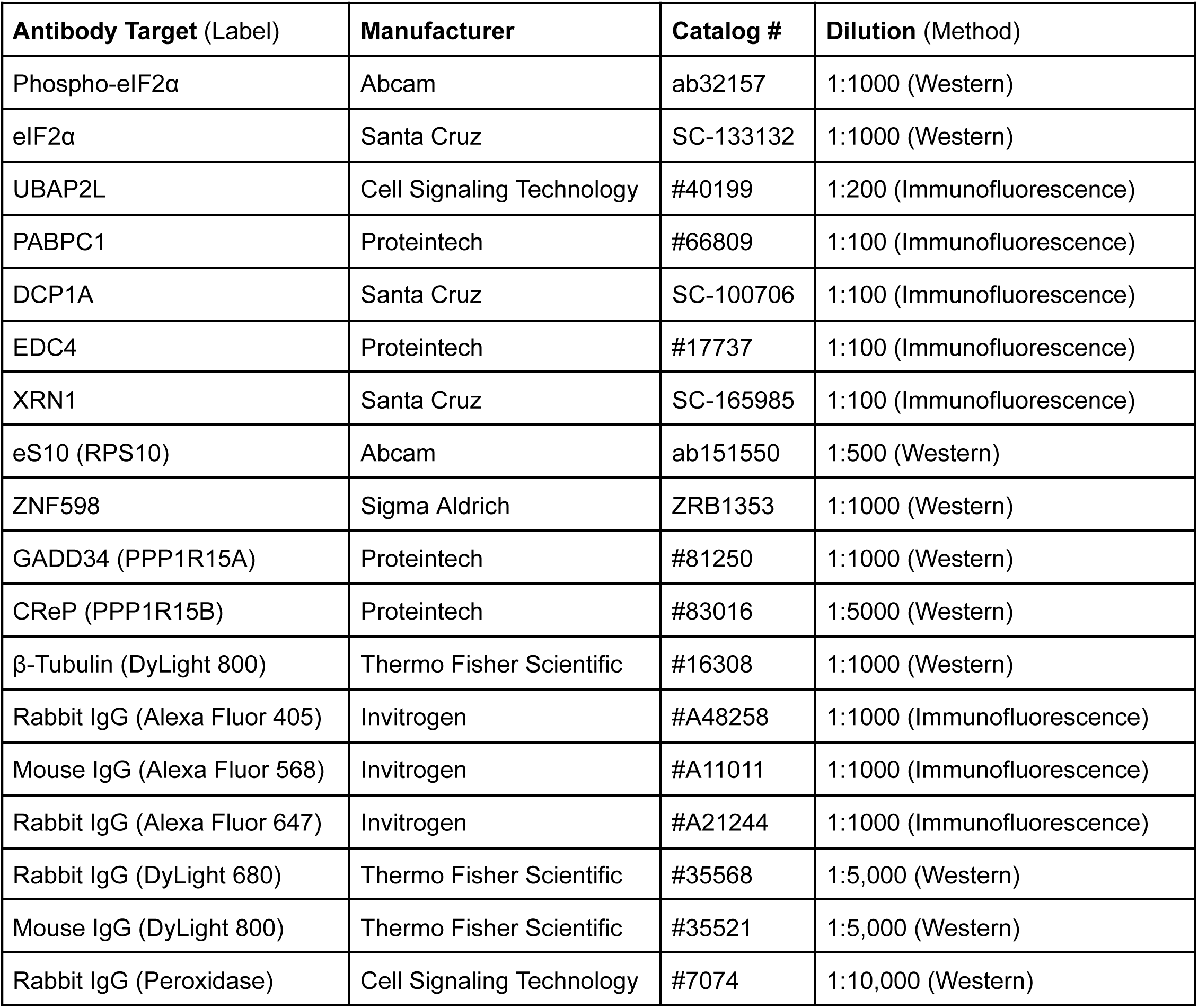
Antibodies used in this study.

Simultaneous immunofluorescence and fluorescence *in situ* hybridization was done as previously described (*52*). Briefly, cells were fixed in 4% paraformaldehyde in PBS for 10 minutes, followed by permeabilization in 0.1% Triton X-100 with 0.2 U/μL of Ribolock RNase inhibitor (Thermo Scientific). Cells were then incubated with primary antibodies (**Table 1**) diluted in PBS with 0.2 U/μL of Ribolock at room temperature for 1 hour, washed three times for 5 minutes each with PBS, and then incubated for 45 minutes at room temperature with fluorescent secondary antibodies (**Table 1**). This was followed by washing three times for 5 minutes in PBS and a second fixation in 4% paraformaldehyde in PBS for 10 minutes. Cells were then incubated in Wash Buffer A (10% formamide in nuclease-free 2x SSC buffer) for 5 minutes followed by hybridization buffer (10% dextran sulfate, 10% formamide in 2x SSC) with Cy3-labeled oligo(dT) (Integrated DNA Technologies - IDT) at 37°C for 20 hours. Hybridization buffer was removed, and each well incubated with Wash Buffer A with NucBlue for 30 minutes at 37°C, followed by an additional 30 minutes in Wash Buffer A. Finally, Wash Buffer A was replaced with Wash Buffer B (2x SSC only) before imaging.

Images of fixed cells were collected on an EVOS M5000 fluorescence microscope (Invitrogen) using a 40X objective. For live cell microscopy, media was exchanged to FluoroBrite (Gibco #A1896701) supplemented with FBE and Glutamax 1 hour before treatment. Imaging was performed using Nikon Elements software and the 40X objective of a Nikon Eclipse Ti2 microscope equipped with a Lumencor Spectra III light engine, Andor Life 888 EMCCD camera, and a Tokai Hit stage top incubator system set to 5% CO_2_ and 37°C. To assess whether stress granules formed over time upon halofuginone treatment, images were collected every 2 hours for 16 hours. To determine if stress granules form in halofuginone-treated cells in the presence or absence of arsenite or rocaglamide A, images were collected every 30 minutes for 3 hours. The percentage of cells with either stress granules and/or P-bodies was quantified manually using the Cell Counter plugin in ImageJ (*169*). Counts were performed from two frames per condition per independent experimental replicate, with n = 3 replicates for every experiment performed.

### Western Blotting

Cells were treated as indicated, washed in PBS, and then 1X RIPA buffer with Halt protease and phosphatase inhibitor cocktail (Fisher Scientific) and benzonase (Fisher) was added directly to the plate. Lysates were removed via scraping, supplemented with 4X Laemmli loading buffer and 10X Bolt Reducing agent (Fisher), heated at 70°C for 10 min, and then loaded onto a Bolt 4-12% Bis-Tris gel (Invitrogen) and electrophoresed in Tris-MES-SDS running buffer (VWR). Proteins were transferred to a PVDF membrane, and the membrane stained with Total Protein Q reagent (Azure Biosystems) and imaged. The membrane was then blocked in TBST buffer with 5% nonfat dry milk for 30 minutes, and incubated with primary antibodies in TBST with milk for 1 hour at room temperature or 16-20 hours at 4°C. The membrane was washed three times for 5 minutes each in TBST, followed by 45 minutes at room temperature with secondary antibodies diluted in TBST buffer with 5% nonfat dry milk. Antibodies used and dilutions are listed in **Table 1**. The membrane was then washed three times for 10 minutes each in TBST. All images were collected with an Azure Biosystems c600 Imaging System, and band intensities were quantified using ImageJ gel analysis tool (*169*). Relative protein levels were determined by normalizing bands to total protein signal in each lane and relative levels of phosphorylated eIF2α were normalized to total eIF2α as indicated in each figure.

### Polysome Profiling

A modified version of the protocol from (*65*) was used for polysome profiling. Briefly, after treatment a 15 cm dish of U-2 OS cells was treated with 100 µg/mL cycloheximide for 10 minutes. Media was aspirated and cells were then washed in ice cold PBS with 100 µg/mL cycloheximide. Cells were then scraped into ice cold lysis buffer containing 20 mM Tris pH 7.5, 150 mM NaCl, 5 mM MgCl_2_, 1% Triton X-100, 1 mM DTT, 1 mg/mL cycloheximide, 80 U/mL Ribolock RNase inhibitor (Thermo Scientific), and 50 U/mL DNase I (Zymo Research). After lysis on ice for 10 minutes, lysates were centrifuged at 20,000 x g for 30 minutes at 4°C to remove cell debris, and then snap frozen in ethanol-dry ice and stored at −80°C. For separation of polysomes, lysates were thawed on ice and loaded onto a 10-50% sucrose gradient in 20 mM Tris pH 7.5, 150 mM KCl and 5 mM MgCl_2_. Gradients were spun at 36,000 RPM for 3 hours at 4°C, and polysome profiles collected using a Biocomp Piston Gradient Fractionator. Absorbance at 260 nm was normalized to 1 for the highest point in each profile. Data were graphed in GraphPad Prism 10.4.0 (GraphPad Software, Boston, Massachusetts USA), exported as .tif files, and the area under the curve measured using the ImageJ wand tracing tool (*169*). The peaks after the free RNP fraction were defined in order as 40S, 60S, and 80S, and all peaks to the right of the 80S were considered as polysomes.

### Bioorthogonal Non-canonical Amino Acid Tagging (BONCAT)

Cells were treated with stressors as indicated, and 10 minutes prior to collection were switched to methionine-free DMEM with either 4 mM methionine as control or 4 mM azidohomoalanine (AHA) for bioorthogonal non-canonical amino acid tagging (BONCAT). Cells were then lysed in 1X RIPA buffer with Halt protease and phosphatase inhibitor cocktail and benzonase, and the Click-&-Go kit (Vector Laboratories) with Alexa 488 alkyne (Invitrogen) used for fluorescent labeling of nascent proteins. The purified labeled protein was supplemented with 4X Laemmli loading buffer and 10X Bolt Reducing agent (Fisher), heated at 70°C for 10 min, and then loaded onto a Bolt 4-12% Bis-Tris gel (Invitrogen) and electrophoresed in Tris-MES-SDS running buffer (VWR). Proteins were transferred to a PVDF membrane, and the clicked fluorophore imaged on the membrane using an Azure Biosystems c600 Imaging System. Total protein staining was then performed with Total Protein Q reagent (Azure Biosystems). Each lane was quantified with the ImageJ gel analysis tool (*169*), and is reported as a ratio of BONCAT signal to total protein signal with the data normalized to the untreated control as 1 for each replicate.

### Quantification and Statistics

All experiments were performed for n = 3 independent experimental replicates, and representative images of a single replicate are shown for each experiment. Data were graphed with bars as the mean and with error bars of standard error of the mean. Points indicate individual measurements, with matching point colors indicating measurements within a single experimental replicate. Statistical significance was assessed using an ordinary one-way ANOVA followed by Tukey’s multiple comparison test and cutoff of p < 0.05 for significance. Graphs and statistical tests were prepared using GraphPad Prism 10.4.0 (GraphPad Software, Boston, Massachusetts USA).

## Acknowledgements

We thank all members of the Moon lab for feedback and helpful discussions. GFP-G3BP1 U-2 OS cells were generously provided by James Burke and Roy Parker. We thank Anthony Antonellis for helpful discussions and feedback. Figure 6 was created with BioRender.com

## Author Contributions

Conceptualization: MB, BD, SLM; Data curation: MB, BD; Formal Analysis: MB; Funding acquisition: MB, NSH, SLM; Investigation: MB; Methodology: MB, NSH, BD, SLM; Project administration: MB, NSH, BD, SLM; Resources: MB, NSH, BD, SLM; Supervision: SLM; Visualization: MB, SLM; Writing–original draft: MB, SLM; Writing–review and editing: MB, NSH, BD, SLM

## Data and Materials Availability

All data needed to evaluate the conclusions in the paper are present in the paper and/or the Supplementary Materials. Source data are provided with this paper.

## Bibliography

1. M. Costa-Mattioli, P. Walter, The integrated stress response: From mechanism to disease. Science 368 (2020).

2. K. Pakos-Zebrucka, I. Koryga, K. Mnich, M. Ljujic, A. Samali, A. M. Gorman, The integrated stress response. EMBO Rep. 17, 1374–1395 (2016).

3. D. W. Sanders, N. Kedersha, D. S. W. Lee, A. R. Strom, V. Drake, J. A. Riback, D. Bracha, J. M. Eeftens, A. Iwanicki, A. Wang, M.-T. Wei, G. Whitney, S. M. Lyons, P. Anderson, W. M. Jacobs, P. Ivanov, C. P. Brangwynne, Competing Protein-RNA Interaction Networks Control Multiphase Intracellular Organization. Cell 181, 306–324.e28 (2020).

4. M. Baymiller, S. Moon, Stress granules as causes and consequences of translation suppression. Antioxid. Redox Signal., doi: 10.1089/ars.2022.0164 (2023).

5. J. Guillén-Boixet, A. Kopach, A. S. Holehouse, S. Wittmann, M. Jahnel, R. Schlüßler, K. Kim, I. R. E. A. Trussina, J. Wang, D. Mateju, I. Poser, S. Maharana, M. Ruer-Gruß, D. Richter, X. Zhang, Y.-T. Chang, J. Guck, A. Honigmann, J. Mahamid, A. A. Hyman, R. V. Pappu, S. Alberti, T. M. Franzmann, RNA-Induced Conformational Switching and Clustering of G3BP Drive Stress Granule Assembly by Condensation. Cell 181, 346–361.e17 (2020).

6. P. Yang, C. Mathieu, R.-M. Kolaitis, P. Zhang, J. Messing, U. Yurtsever, Z. Yang, J. Wu, Y. Li, Q. Pan, J. Yu, E. W. Martin, T. Mittag, H. J. Kim, J. P. Taylor, G3BP1 Is a Tunable Switch that Triggers Phase Separation to Assemble Stress Granules. Cell 181, 325–345.e28 (2020).

7. S. L. Moon, T. Morisaki, A. Khong, K. Lyon, R. Parker, T. J. Stasevich, Multicolour single-molecule tracking of mRNA interactions with RNP granules. Nat. Cell Biol. 21, 162–168 (2019).

8. J. H. Wilbertz, F. Voigt, I. Horvathova, G. Roth, Y. Zhan, J. A. Chao, Single-Molecule Imaging of mRNA Localization and Regulation during the Integrated Stress Response. Mol. Cell 73, 946–958.e7 (2019).

9. S. Pitchiaya, M. D. A. Mourao, A. P. Jalihal, L. Xiao, X. Jiang, A. M. Chinnaiyan, S. Schnell, N. G. Walter, Dynamic Recruitment of Single RNAs to Processing Bodies Depends on RNA Functionality. Mol. Cell 74, 521–533.e6 (2019).

10. J. Zhang, K. Okabe, T. Tani, T. Funatsu, Dynamic association-dissociation and harboring of endogenous mRNAs in stress granules. J. Cell Sci. 124, 4087–4095 (2011).

11. S. Mollet, N. Cougot, A. Wilczynska, F. Dautry, M. Kress, E. Bertrand, D. Weil, Translationally repressed mRNA transiently cycles through stress granules during stress. Mol. Biol. Cell 19, 4469–4479 (2008).

12. N. Kedersha, G. Stoecklin, M. Ayodele, P. Yacono, J. Lykke-Andersen, M. J. Fritzler, D. Scheuner, R. J. Kaufman, D. E. Golan, P. Anderson, Stress granules and processing bodies are dynamically linked sites of mRNP remodeling. J. Cell Biol. 169, 871–884 (2005).

13. N. Kedersha, M. R. Cho, W. Li, P. W. Yacono, S. Chen, N. Gilks, D. E. Golan, P. Anderson, Dynamic shuttling of TIA-1 accompanies the recruitment of mRNA to mammalian stress granules. J. Cell Biol. 151, 1257–1268 (2000).

14. A. Khong, T. Matheny, S. Jain, S. F. Mitchell, J. R. Wheeler, R. Parker, The Stress Granule Transcriptome Reveals Principles of mRNA Accumulation in Stress Granules. Mol. Cell 68, 808–820.e5 (2017).

15. B. Van Treeck, D. S. W. Protter, T. Matheny, A. Khong, C. D. Link, R. Parker, RNA self-assembly contributes to stress granule formation and defining the stress granule transcriptome. Proc. Natl. Acad. Sci. U. S. A. 115, 2734–2739 (2018).

16. T. Matheny, B. Van Treeck, T. N. Huynh, R. Parker, RNA partitioning into stress granules is based on the summation of multiple interactions. RNA 27, 174–189 (2021).

17. Z. Ren, W. Tang, L. Peng, P. Zou, Profiling stress-triggered RNA condensation with photocatalytic proximity labeling. Nat. Commun. 14, 7390 (2023).

18. S. Namkoong, A. Ho, Y. M. Woo, H. Kwak, J. H. Lee, Systematic Characterization of Stress-Induced RNA Granulation. Mol. Cell 70, 175–187.e8 (2018).

19. Matheny Tyler, Rao Bhalchandra S., Parker Roy, Transcriptome-Wide Comparison of Stress Granules and P-Bodies Reveals that Translation Plays a Major Role in RNA Partitioning. Mol. Cell. Biol. 39, e00313–19 (2019).

20. A. Padrón, S. Iwasaki, N. T. Ingolia, Proximity RNA Labeling by APEX-Seq Reveals the Organization of Translation Initiation Complexes and Repressive RNA Granules. Mol. Cell 75, 875–887.e5 (2019).

21. A. Hubstenberger, M. Courel, M. Bénard, S. Souquere, M. Ernoult-Lange, R. Chouaib, Z. Yi, J.-B. Morlot, A. Munier, M. Fradet, M. Daunesse, E. Bertrand, G. Pierron, J. Mozziconacci, M. Kress, D. Weil, P-body purification reveals the condensation of repressed mRNA regulons. Mol. Cell 68, 144–157.e5 (2017).

22. J. M. Burke, O. C. Ratnayake, J. M. Watkins, R. Perera, R. Parker, G3BP1-dependent condensation of translationally inactive viral RNAs antagonizes infection. Sci Adv 10, eadk8152 (2024).

23. C. Desroches Altamirano, M.-K. Kang, M. A. Jordan, T. Borianne, I. Dilmen, M. Gnädig, A. von Appen, A. Honigmann, T. M. Franzmann, S. Alberti, eIF4F is a thermo-sensing regulatory node in the translational heat shock response. Mol. Cell, doi: 10.1016/j.molcel.2024.02.038 (2024).

24. J. A. Riback, C. D. Katanski, J. L. Kear-Scott, E. V. Pilipenko, A. E. Rojek, T. R. Sosnick, D. A. Drummond, Stress-Triggered Phase Separation Is an Adaptive, Evolutionarily Tuned Response. Cell 168, 1028–1040.e19 (2017).

25. A. Dubinski, M. Gagné, S. Peyrard, D. Gordon, K. Talbot, C. Vande Velde, Stress granule assembly in vivo is deficient in the CNS of mutant TDP-43 ALS mice. Hum. Mol. Genet. (2022).

26. B. Di Stefano, E.-C. Luo, C. Haggerty, S. Aigner, J. Charlton, J. Brumbaugh, F. Ji, I. Rabano Jiménez, K. J. Clowers, A. J. Huebner, K. Clement, I. Lipchina, M. A. C. de Kort, A. Anselmo, J. Pulice, M. F. M. Gerli, H. Gu, S. P. Gygi, R. I. Sadreyev, A. Meissner, G. W. Yeo, K. Hochedlinger, The RNA helicase DDX6 controls cellular plasticity by modulating P-body homeostasis. Cell Stem Cell 25, 622–638.e13 (2019).

27. L. C. Reineke, S. A. Cheema, J. Dubrulle, J. R. Neilson, Chronic starvation induces noncanonical pro-death stress granules. J. Cell Sci. 131 (2018).

28. Y. R. Li, O. D. King, J. Shorter, A. D. Gitler, Stress granules as crucibles of ALS pathogenesis. J. Cell Biol. 201, 361–372 (2013).

29. T. Wang, X. Tian, H. B. Kim, Y. Jang, Z. Huang, C. H. Na, J. Wang, Intracellular energy controls dynamics of stress-induced ribonucleoprotein granules. Nat. Commun. 13, 5584 (2022).

30. S. Kodali, L. Proietti, G. Valcarcel, A. V. López-Rubio, P. Pessina, T. Eder, J. Shi, A. Jen, N. Lupión-Garcia, A. C. Starner, M. D. Bartels, Y. Cui, C. M. Sands, A. Planas-Riverola, A. Martínez, T. Velasco-Hernandez, L. Tomás-Daza, B. Alber, G. Manhart, I. M. Mayer, K. Kollmann, A. Fatica, P. Menendez, E. Shishkova, R. E. Rau, B. M. Javierre, J. Coon, Q. Chen, E. L. Van Nostrand, J. L. Sardina, F. Grebien, B. Di Stefano, RNA sequestration in P-bodies sustains myeloid leukaemia. Nat. Cell Biol. 26, 1745–1758 (2024).

31. N. J. Boon, R. A. Oliveira, P.-R. Körner, A. Kochavi, S. Mertens, Y. Malka, R. Voogd, S. E. M. van der Horst, M. A. Huismans, L. P. Smabers, J. M. Draper, L. F. A. Wessels, P. Haahr, J. M. L. Roodhart, T. N. M. Schumacher, H. J. Snippert, R. Agami, T. R. Brummelkamp, DNA damage induces p53-independent apoptosis through ribosome stalling. Science 384, 785–792 (2024).

32. M. Stoneley, R. F. Harvey, T. E. Mulroney, R. Mordue, R. Jukes-Jones, K. Cain, K. S. Lilley, R. Sawarkar, E. Willis, Unresolved stalled ribosome complexes restrict cell-cycle progression after genotoxic stress. Mol. Cell 82, 1557–1572.e7 (2022).

33. G. Snieckute, A. V. Genzor, A. C. Vind, L. Ryder, M. Stoneley, S. Chamois, R. Dreos, C. Nordgaard, F. Sass, M. Blasius, A. R. López, S. H. Brynjólfsdóttir, K. L. Andersen, A. E. Willis, L. B. Frankel, S. S. Poulsen, D. Gatfield, Z. Gerhart-Hines, C. Clemmensen, S. Bekker-Jensen, Ribosome stalling is a signal for metabolic regulation by the ribotoxic stress response. Cell Metab. 34, 2036–2046.e8 (2022).

34. L. L. Yan, H. S. Zaher, Ribosome quality control antagonizes the activation of the integrated stress response on colliding ribosomes. Mol. Cell 81, 614–628.e4 (2021).

35. L. Ryder, F. S. Arendrup, J. F. Martínez, G. Snieckute, C. Pecorari, R. A. Shah, A. H. Lund, M. Blasius, S. Bekker-Jensen, Nitric oxide-induced ribosome collision activates ribosomal surveillance mechanisms. Cell Death Dis. 14, 467 (2023).

36. T. L. Keller, D. Zocco, M. S. Sundrud, M. Hendrick, M. Edenius, J. Yum, Y.-J. Kim, H.-K. Lee, J. F. Cortese, D. F. Wirth, J. D. Dignam, A. Rao, C.-Y. Yeo, R. Mazitschek, M. Whitman, Halofuginone and other febrifugine derivatives inhibit prolyl-tRNA synthetase. Nat. Chem. Biol. 8, 311–317 (2012).

37. M. S. Sundrud, S. B. Koralov, M. Feuerer, D. P. Calado, A. E. Kozhaya, A. Rhule-Smith, R. E. Lefebvre, D. Unutmaz, R. Mazitschek, H. Waldner, M. Whitman, T. Keller, A. Rao, Halofuginone inhibits TH17 cell differentiation by activating the amino acid starvation response. Science 324, 1334–1338 (2009).

38. J. Misra, K. R. Carlson, D. F. Spandau, R. C. Wek, Multiple mechanisms activate GCN2 eIF2 kinase in response to diverse stress conditions. Nucleic Acids Res., doi: 10.1093/nar/gkae006 (2024).

39. J. Dong, H. Qiu, M. Garcia-Barrio, J. Anderson, A. G. Hinnebusch, Uncharged tRNA activates GCN2 by displacing the protein kinase moiety from a bipartite tRNA-binding domain. Mol. Cell 6, 269–279 (2000).

40. J. Z. Yin, A. F. A. Keszei, S. Houliston, F. Filandr, J. Beenstock, S. Daou, J. Kitaygorodsky, D. C. Schriemer, M. T. Mazhab-Jafari, A.-C. Gingras, F. Sicheri, The HisRS-like domain of GCN2 is a pseudoenzyme that can bind uncharged tRNA. Structure 32, 795–811.e6 (2024).

41. S. A. Wek, S. Zhu, R. C. Wek, The histidyl-tRNA synthetase-related sequence in the eIF-2 alpha protein kinase GCN2 interacts with tRNA and is required for activation in response to starvation for different amino acids. Mol. Cell. Biol. 15, 4497–4506 (1995).

42. H. P. Harding, A. Ordonez, F. Allen, L. Parts, A. J. Inglis, R. L. Williams, D. Ron, The ribosomal P-stalk couples amino acid starvation to GCN2 activation in mammalian cells. Elife 8 (2019).

43. A. J. Inglis, G. R. Masson, S. Shao, O. Perisic, S. H. McLaughlin, R. S. Hegde, R. L. Williams, Activation of GCN2 by the ribosomal P-stalk. Proc. Natl. Acad. Sci. U. S. A. 116, 4946–4954 (2019).

44. C. C.-C. Wu, A. Peterson, B. Zinshteyn, S. Regot, R. Green, Ribosome Collisions Trigger General Stress Responses to Regulate Cell Fate. Cell 182, 404–416.e14 (2020).

45. R. Ishimura, G. Nagy, I. Dotu, J. H. Chuang, S. L. Ackerman, Activation of GCN2 kinase by ribosome stalling links translation elongation with translation initiation. Elife 5 (2016).

46. G. R. Masson, Towards a model of GCN2 activation. Biochem. Soc. Trans. 47, 1481–1488 (2019).

47. A. A. Pochopien, B. Beckert, S. Kasvandik, O. Berninghausen, R. Beckmann, T. Tenson, D. N. Wilson, Structure of Gcn1 bound to stalled and colliding 80S ribosomes. Proc. Natl. Acad. Sci. U. S. A. 118 (2021).

48. N. S. Helton, B. Dodd, S. L. Moon, Stress-Induced Gene mRNA Condensation and Expression are Governed by Ribosome Association and the Stress Granule Proteins G3BP1/2, Social Science Research Network (2024). https://papers.ssrn.com/abstract=4913224.

49. A. Khong, R. Parker, mRNP architecture in translating and stress conditions reveals an ordered pathway of mRNP compaction. J. Cell Biol. 217, 4124–4140 (2018).

50. D. Teixeira, U. Sheth, M. A. Valencia-Sanchez, M. Brengues, R. Parker, Processing bodies require RNA for assembly and contain nontranslating mRNAs. RNA 11, 371–382 (2005).

51. R. Mazroui, R. Sukarieh, M.-E. Bordeleau, R. J. Kaufman, P. Northcote, J. Tanaka, I. Gallouzi, J. Pelletier, Inhibition of Ribosome Recruitment Induces Stress Granule Formation Independently of Eukaryotic Initiation Factor 2α Phosphorylation. MBoC 17, 4212–4219 (2006).

52. S. L. Moon, T. Morisaki, T. J. Stasevich, R. Parker, Coupling of translation quality control and mRNA targeting to stress granules. J. Cell Biol. 219 (2020).

53. O. Brandman, J. Stewart-Ornstein, D. Wong, A. Larson, C. C. Williams, G.-W. Li, S. Zhou, D. King, P. S. Shen, J. Weibezahn, J. G. Dunn, S. Rouskin, T. Inada, A. Frost, J. S. Weissman, A ribosome-bound quality control complex triggers degradation of nascent peptides and signals translation stress. Cell 151, 1042–1054 (2012).

54. S. Juszkiewicz, S. H. Speldewinde, L. Wan, J. Q. Svejstrup, R. S. Hegde, The ASC-1 Complex Disassembles Collided Ribosomes. Mol. Cell 79, 603–614.e8 (2020).

55. M. K. Doma, R. Parker, Endonucleolytic cleavage of eukaryotic mRNAs with stalls in translation elongation. Nature 440, 561–564 (2006).

56. C. J. Shoemaker, D. E. Eyler, R. Green, Dom34:Hbs1 promotes subunit dissociation and peptidyl-tRNA drop-off to initiate no-go decay. Science 330, 369–372 (2010).

57. M. Terrey, S. I. Adamson, A. L. Gibson, T. Deng, R. Ishimura, J. H. Chuang, S. L. Ackerman, GTPBP1 resolves paused ribosomes to maintain neuronal homeostasis. Elife 10 (2021).

58. R. Ishimura, G. Nagy, I. Dotu, H. Zhou, X.-L. Yang, P. Schimmel, S. Senju, Y. Nishimura, J. H. Chuang, S. L. Ackerman, RNA function. Ribosome stalling induced by mutation of a CNS-specific tRNA causes neurodegeneration. Science 345, 455–459 (2014).

59. A. C. Vind, G. Snieckute, M. Blasius, C. Tiedje, N. Krogh, D. B. Bekker-Jensen, K. L. Andersen, C. Nordgaard, M. A. X. Tollenaere, A. H. Lund, J. V. Olsen, H. Nielsen, S. Bekker-Jensen, ZAKα Recognizes Stalled Ribosomes through Partially Redundant Sensor Domains. Mol. Cell 78, 700–713.e7 (2020).

60. A. S. Rahmanto, C. J. Blum, C. Scalera, J. B. Heidelberger, M. Mesitov, D. Horn-Ghetko, J. F. Gräf, I. Mikicic, R. Hobrecht, A. Orekhova, M. Ostermaier, S. Ebersberger, M. M. Möckel, N. Krapoth, N. Da Silva Fernandes, A. Mizi, Y. Zhu, J.-X. Chen, C. Choudhary, A. Papantonis, H. D. Ulrich, B. A. Schulman, J. König, P. Beli, K6-linked ubiquitylation marks formaldehyde-induced RNA-protein crosslinks for resolution. Mol. Cell 83, 4272–4289.e10 (2023).

61. S. Zhao, J. Cordes, K. M. Caban, M. J. Götz, T. Mackens-Kiani, A. J. Veltri, N. K. Sinha, P. Weickert, S. Kaya, G. Hewitt, D. D. Nedialkova, T. Fröhlich, R. Beckmann, A. R. Buskirk, R. Green, J. Stingele, RNF14-dependent atypical ubiquitylation promotes translation-coupled resolution of RNA-protein crosslinks. Mol. Cell 83, 4290–4303.e9 (2023).

62. J. Misra, M. J. Holmes, E. T Mirek, M. Langevin, H.-G. Kim, K. R. Carlson, M. Watford, X. C. Dong, T. G. Anthony, R. C. Wek, Discordant regulation of eIF2 kinase GCN2 and mTORC1 during nutrient stress. Nucleic Acids Res. 49, 5726–5742 (2021).

63. Y. Kim, M. S. Sundrud, C. Zhou, M. Edenius, D. Zocco, K. Powers, M. Zhang, R. Mazitschek, A. Rao, C.-Y. Yeo, E. H. Noss, M. B. Brenner, M. Whitman, T. L. Keller, Aminoacyl-tRNA synthetase inhibition activates a pathway that branches from the canonical amino acid response in mammalian cells. Proc. Natl. Acad. Sci. U. S. A. 117, 8900–8911 (2020).

64. A. P. Pitera, M. Szaruga, S.-Y. Peak-Chew, S. W. Wingett, A. Bertolotti, Cellular responses to halofuginone reveal a vulnerability of the GCN2 branch of the integrated stress response. EMBO J. 41, e109985 (2022).

65. L. Ferguson, H. E. Upton, S. C. Pimentel, A. Mok, L. F. Lareau, K. Collins, N. T. Ingolia, Streamlined and sensitive mono- and di-ribosome profiling in yeast and human cells. Nat. Methods 20, 1704–1715 (2023).

66. R. Muller, Z. A. Meacham, L. Ferguson, N. T. Ingolia, CiBER-seq dissects genetic networks by quantitative CRISPRi profiling of expression phenotypes. Science 370 (2020).

67. S. Battu, S. Afroz, J. Giddaluru, S. Naz, W. Huang, S. S. Khumukcham, R. A. Khan, S. Y. Bhat, I. A. Qureshi, B. Manavathi, A. A. Khan, A. August, S. E. Hasnain, N. Khan, Amino acid starvation sensing dampens IL-1β production by activating riboclustering and autophagy. PLoS Biol. 16, e2005317 (2018).

68. M. Pines, A. Nagler, Halofuginone: a novel antifibrotic therapy. Gen. Pharmacol. 30, 445–450 (1998).

69. M. Elkin, I. Ariel, H. Q. Miao, A. Nagler, M. Pines, N. de-Groot, A. Hochberg, I. Vlodavsky, Inhibition of bladder carcinoma angiogenesis, stromal support, and tumor growth by halofuginone. Cancer Res. 59, 4111–4118 (1999).

70. D.-K. Lee, S. H. Jo, E. S. Lee, K. B. Ha, N. W. Park, D.-H. Kong, S.-I. Park, J. S. Park, C. H. Chung, DWN12088, A Prolyl-tRNA Synthetase Inhibitor, Alleviates Hepatic Injury in Nonalcoholic Steatohepatitis. Diabetes Metab. J., doi: 10.4093/dmj.2022.0367 (2024).

71. S. Rai, M. Szaruga, A. P. Pitera, A. Bertolotti, Integrated stress response activator halofuginone protects mice from diabetes-like phenotypes. J. Cell Biol. 223 (2024).

72. A. Antonellis, R. E. Ellsworth, N. Sambuughin, I. Puls, A. Abel, S.-Q. Lee-Lin, A. Jordanova, I. Kremensky, K. Christodoulou, L. T. Middleton, K. Sivakumar, V. Ionasescu, B. Funalot, J. M. Vance, L. G. Goldfarb, K. H. Fischbeck, E. D. Green, Glycyl tRNA synthetase mutations in Charcot-Marie-Tooth disease type 2D and distal spinal muscular atrophy type V. Am. J. Hum. Genet. 72, 1293–1299 (2003).

73. A. Jordanova, J. Irobi, F. P. Thomas, P. Van Dijck, K. Meerschaert, M. Dewil, I. Dierick, A. Jacobs, E. De Vriendt, V. Guergueltcheva, C. V. Rao, I. Tournev, F. A. A. Gondim, M. D’Hooghe, V. Van Gerwen, P. Callaerts, L. Van Den Bosch, J.-P. Timmermans, W. Robberecht, J. Gettemans, J. M. Thevelein, P. De Jonghe, I. Kremensky, V. Timmerman, Disrupted function and axonal distribution of mutant tyrosyl-tRNA synthetase in dominant intermediate Charcot-Marie-Tooth neuropathy. Nat. Genet. 38, 197–202 (2006).

74. P. Latour, C. Thauvin-Robinet, C. Baudelet-Méry, P. Soichot, V. Cusin, L. Faivre, M.-C. Locatelli, M. Mayençon, A. Sarcey, E. Broussolle, W. Camu, A. David, R. Rousson, A major determinant for binding and aminoacylation of tRNA(Ala) in cytoplasmic Alanyl-tRNA synthetase is mutated in dominant axonal Charcot-Marie-Tooth disease. Am. J. Hum. Genet. 86, 77–82 (2010).

75. P.-C. Tsai, B.-W. Soong, I. Mademan, Y.-H. Huang, C.-R. Liu, C.-T. Hsiao, H.-T. Wu, T.-T. Liu, Y.-T. Liu, Y.-T. Tseng, K.-P. Lin, U.-C. Yang, K. W. Chung, B.-O. Choi, G. A. Nicholson, M. L. Kennerson, C.-C. Chan, P. De Jonghe, T.-H. Cheng, Y.-C. Liao, S. Züchner, J. Baets, Y.-C. Lee, A recurrent WARS mutation is a novel cause of autosomal dominant distal hereditary motor neuropathy. Brain 140, 1252–1266 (2017).

76. A. Vester, G. Velez-Ruiz, H. M. McLaughlin, NISC Comparative Sequencing Program, J. R. Lupski, K. Talbot, J. M. Vance, S. Züchner, R. H. Roda, K. H. Fischbeck, L. G. Biesecker, G. Nicholson, A. A. Beg, A. Antonellis, A loss-of-function variant in the human histidyl-tRNA synthetase (HARS) gene is neurotoxic in vivo. Hum. Mutat. 34, 191–199 (2013).

77. D. Beijer, S. Marte, J. C. Li, W. De Ridder, J. Z. Chen, A. L. D. Tadenev, K. E. Miers, T. Deconinck, R. Macdonell, W. Marques Jr, P. De Jonghe, S. L. Pratt, R. Meyer-Schuman, S. Züchner, A. Antonellis, R. W. Burgess, J. Baets, Dominant NARS1 mutations causing axonal Charcot-Marie-Tooth disease expand NARS1-associated diseases. Brain Commun. 6, fcae070 (2024).

78. M. E. Kuo, A. Antonellis, Ubiquitously Expressed Proteins and Restricted Phenotypes: Exploring Cell-Specific Sensitivities to Impaired tRNA Charging. Trends Genet. 36, 105–117 (2020).

79. A. K. Turvey, G. A. Horvath, A. R. O. Cavalcanti, Aminoacyl-tRNA synthetases in human health and disease. Front. Physiol. 13, 1029218 (2022).

80. D. Jin, S. A. Wek, N. T. Kudlapur, W. A. Cantara, M. Bakhtina, R. C. Wek, K. Musier-Forsyth, Disease-associated mutations in a bifunctional aminoacyl-tRNA synthetase gene elicit the integrated stress response. J. Biol. Chem. 297, 101203 (2021).

81. D. Jin, S. A. Wek, R. A. Cordova, R. C. Wek, D. Lacombe, V. Michaud, K. Musier-Forsyth, Aminoacylation-defective bi-allelic mutations in human EPRS1 associated with psychomotor developmental delay, epilepsy, and deafness. Clin. Genet. 103, 358–363 (2023).

82. D. Khan, I. Ramachandiran, K. Vasu, A. China, K. Khan, F. Cumbo, D. Halawani, F. Terenzi, I. Zin, B. Long, G. Costain, S. Blaser, A. Carnevale, V. Gogonea, R. Dutta, D. Blankenberg, G. Yoon, P. L. Fox, Homozygous EPRS1 missense variant causing hypomyelinating leukodystrophy-15 alters variant-distal mRNA m6A site accessibility. Nat. Commun. 15, 4284 (2024).

83. A. Zuko, M. Mallik, R. Thompson, E. L. Spaulding, A. R. Wienand, M. Been, A. L. D. Tadenev, N. van Bakel, C. Sijlmans, L. A. Santos, J. Bussmann, M. Catinozzi, S. Das, D. Kulshrestha, R. W. Burgess, Z. Ignatova, E. Storkebaum, tRNA overexpression rescues peripheral neuropathy caused by mutations in tRNA synthetase. Science 373, 1161–1166 (2021).

84. E. L. Spaulding, T. J. Hines, P. Bais, A. L. D. Tadenev, R. Schneider, D. Jewett, B. Pattavina, S. L. Pratt, K. H. Morelli, M. G. Stum, D. P. Hill, C. Gobet, M. Pipis, M. M. Reilly, M. J. Jennings, R. Horvath, Y. Bai, M. E. Shy, B. Alvarez-Castelao, E. M. Schuman, L. P. Bogdanik, E. Storkebaum, R. W. Burgess, The integrated stress response contributes to tRNA synthetase-associated peripheral neuropathy. Science 373, 1156–1161 (2021).

85. D. C. Dieterich, A. J. Link, J. Graumann, D. A. Tirrell, E. M. Schuman, Selective identification of newly synthesized proteins in mammalian cells using bioorthogonal noncanonical amino acid tagging (BONCAT). Proc. Natl. Acad. Sci. U. S. A. 103, 9482–9487 (2006).

86. M. H. Bengtson, C. A. P. Joazeiro, Role of a ribosome-associated E3 ubiquitin ligase in protein quality control. Nature 467, 470–473 (2010).

87. J. M. Burke, E. T. Lester, D. Tauber, R. Parker, RNase L promotes the formation of unique ribonucleoprotein granules distinct from stress granules. J. Biol. Chem. 295, 1426–1438 (2020).

88. R. Parker, U. Sheth, P bodies and the control of mRNA translation and degradation. Mol. Cell 25, 635–646 (2007).

89. U. Sheth, R. Parker, Decapping and decay of messenger RNA occur in cytoplasmic processing bodies. Science 300, 805–808 (2003).

90. M. E. Azzam, I. D. Algranati, Mechanism of puromycin action: fate of ribosomes after release of nascent protein chains from polysomes. Proc. Natl. Acad. Sci. U. S. A. 70, 3866–3869 (1973).

91. S. Juszkiewicz, V. Chandrasekaran, Z. Lin, S. Kraatz, V. Ramakrishnan, R. S. Hegde, ZNF598 Is a Quality Control Sensor of Collided Ribosomes. Mol. Cell 72, 469–481.e7 (2018).

92. M. Narita, T. Denk, Y. Matsuo, T. Sugiyama, C. Kikuguchi, S. Ito, N. Sato, T. Suzuki, S. Hashimoto, I. Machová, P. Tesina, R. Beckmann, T. Inada, A distinct mammalian disome collision interface harbors K63-linked polyubiquitination of uS10 to trigger hRQT-mediated subunit dissociation. Nat. Commun. 13, 6411 (2022).

93. E. Sundaramoorthy, M. Leonard, R. Mak, J. Liao, A. Fulzele, E. J. Bennett, ZNF598 and RACK1 Regulate Mammalian Ribosome-Associated Quality Control Function by Mediating Regulatory 40S Ribosomal Ubiquitylation. Mol. Cell 65, 751–760.e4 (2017).

94. S. Juszkiewicz, R. S. Hegde, Initiation of Quality Control during Poly(A) Translation Requires Site-Specific Ribosome Ubiquitination. Mol. Cell 65, 743–750.e4 (2017).

95. D. H. Goldman, N. M. Livingston, J. Movsik, B. Wu, R. Green, Live-cell imaging reveals kinetic determinants of quality control triggered by ribosome stalling. Mol. Cell 81, 1830–1840.e8 (2021).

96. A. P. Grollman, Z. Jarkovsky, “Emetine and Related Alkaloids” in *Mechanism of Action of Antimicrobial and Antitumor Agents* (Springer Berlin Heidelberg, Berlin, Heidelberg, 1975), pp. 420–435.

97. S. Juszkiewicz, G. Slodkowicz, Z. Lin, P. Freire-Pritchett, S.-Y. Peak-Chew, R. S. Hegde, Ribosome collisions trigger cis-acting feedback inhibition of translation initiation. Elife 9 (2020).

98. S. Lageix, J. Zhang, S. Rothenburg, A. G. Hinnebusch, Interaction between the tRNA-binding and C-terminal domains of Yeast Gcn2 regulates kinase activity in vivo. PLoS Genet. 11, e1004991 (2015).

99. R. C. Wek, B. M. Jackson, A. G. Hinnebusch, Juxtaposition of domains homologous to protein kinases and histidyl-tRNA synthetases in GCN2 protein suggests a mechanism for coupling GCN4 expression to amino acid availability. Proc. Natl. Acad. Sci. U. S. A. 86, 4579–4583 (1989).

100. N. Cougot, S. Babajko, B. Séraphin, Cytoplasmic foci are sites of mRNA decay in human cells. J. Cell Biol. 165, 31–40 (2004).

101. S. Markmiller, S. Soltanieh, K. L. Server, R. Mak, W. Jin, M. Y. Fang, E.-C. Luo, F. Krach, D. Yang, A. Sen, A. Fulzele, J. M. Wozniak, D. J. Gonzalez, M. W. Kankel, F.-B. Gao, E. J. Bennett, E. Lécuyer, G. W. Yeo, Context-Dependent and Disease-Specific Diversity in Protein Interactions within Stress Granules. Cell 172, 590–604.e13 (2018).

102. F. J. Martinez, G. A. Pratt, E. L. Van Nostrand, R. Batra, S. C. Huelga, K. Kapeli, P. Freese, S. J. Chun, K. Ling, C. Gelboin-Burkhart, L. Fijany, H. C. Wang, J. K. Nussbacher, S. M. Broski, H. J. Kim, R. Lardelli, B. Sundararaman, J. P. Donohue, A. Javaherian, J. Lykke-Andersen, S. Finkbeiner, C. F. Bennett, M. Ares Jr, C. B. Burge, J. P. Taylor, F. Rigo, G. W. Yeo, Protein-RNA Networks Regulated by Normal and ALS-Associated Mutant HNRNPA2B1 in the Nervous System. Neuron 92, 780–795 (2016).

103. W. Paetz, G. Nass, Biochemical and immunological characterization of threonyl-tRNA synthetase of two borrelidin-resistant mutants of Escherichia coli K12. Eur. J. Biochem. 35, 331–337 (1973).

104. A. Sidhu, J. R. Miller, A. Tripathi, D. M. Garshott, A. L. Brownell, D. J. Chiego, C. Arevang, Q. Zeng, L. C. Jackson, S. A. Bechler, M. U. Callaghan, G. H. Yoo, S. Sethi, H.-S. Lin, J. H. Callaghan, G. Tamayo-Castillo, D. H. Sherman, R. J. Kaufman, A. M. Fribley, Borrelidin Induces the Unfolded Protein Response in Oral Cancer Cells and Chop-Dependent Apoptosis. ACS Med. Chem. Lett. 6, 1122–1127 (2015).

105. K. N. D’Orazio, R. Green, Ribosome states signal RNA quality control. Mol. Cell 81, 1372–1383 (2021).

106. M. C. J. Yip, S. Shao, Detecting and rescuing stalled ribosomes. Trends Biochem. Sci. 46, 731–743 (2021).

107. H. Sadlish, G. Galicia-Vazquez, C. G. Paris, T. Aust, B. Bhullar, L. Chang, S. B. Helliwell, D. Hoepfner, B. Knapp, R. Riedl, S. Roggo, S. Schuierer, C. Studer, J. A. Porco Jr, J. Pelletier, N. R. Movva, Evidence for a functionally relevant rocaglamide binding site on the eIF4A-RNA complex. ACS Chem. Biol. 8, 1519–1527 (2013).

108. K. L. Wollen, L. Hagen, C. B. Vågbø, R. Rabe, T. S. Iveland, P. A. Aas, A. Sharma, B. Sporsheim, H. O. Erlandsen, V. Palibrk, M. Bjørås, D. M. Fonseca, N. Mosammaparast, G. Slupphaug, ALKBH3 partner ASCC3 mediates P-body formation and selective clearance of MMS-induced 1-methyladenosine and 3-methylcytosine from mRNA. J. Transl. Med. 19, 287 (2021).

109. K. Arimoto, H. Fukuda, S. Imajoh-Ohmi, H. Saito, M. Takekawa, Formation of stress granules inhibits apoptosis by suppressing stress-responsive MAPK pathways. Nat. Cell Biol. 10, 1324–1332 (2008).

110. A. Aulas, S. M. Lyons, M. M. Fay, P. Anderson, P. Ivanov, Nitric oxide triggers the assembly of “type II” stress granules linked to decreased cell viability. Cell Death Dis. 9, 1–14 (2018).

111. P. Pietras, A. Aulas, M. M. Fay, M. Leśniczak-Staszak, M. Sowiński, S. M. Lyons, W. Szaflarski, P. Ivanov, Translation inhibition and suppression of stress granules formation by cisplatin. Biomed. Pharmacother. 145, 112382 (2022).

112. J. L. Martin, S. J. Terry, J. E. Gale, S. J. Dawson, The ototoxic drug cisplatin localises to stress granules altering their dynamics and composition. J. Cell Sci., doi: 10.1242/jcs.260590 (2023).

113. N. K. Sinha, C. McKenney, Z. Y. Yeow, J. J. Li, K. H. Nam, T. M. Yaron-Barir, J. L. Johnson, E. M. Huntsman, L. C. Cantley, A. Ordureau, S. Regot, R. Green, The ribotoxic stress response drives UV-mediated cell death. Cell, doi: 10.1016/j.cell.2024.05.018 (2024).

114. S. Ying, D. A. Khaperskyy, UV damage induces G3BP1-dependent stress granule formation that is not driven by mTOR inhibition-mediated translation arrest. J. Cell Sci. 133 (2020).

115. A. Aulas, M. M. Fay, S. M. Lyons, C. A. Achorn, N. Kedersha, P. Anderson, P. Ivanov, Stress-specific differences in assembly and composition of stress granules and related foci. J. Cell Sci. 130, 927–937 (2017).

116. Y. Zhou, A. Panhale, M. Shvedunova, M. Balan, A. Gomez-Auli, H. Holz, J. Seyfferth, M. Helmstädter, S. Kayser, Y. Zhao, N. U. Erdogdu, I. Grzadzielewska, G. Mittler, T. Manke, A. Akhtar, RNA damage compartmentalization by DHX9 stress granules. Cell, doi: 10.1016/j.cell.2024.02.028 (2024).

117. K. Best, K. Ikeuchi, L. Kater, D. Best, J. Musial, Y. Matsuo, O. Berninghausen, T. Becker, T. Inada, R. Beckmann, Structural basis for clearing of ribosome collisions by the RQT complex. Nat. Commun. 14, 921 (2023).

118. A. A. Morgan, E. Rubenstein, Proline: the distribution, frequency, positioning, and common functional roles of proline and polyproline sequences in the human proteome. PLoS One 8, e53785 (2013).

119. A. J. Veltri, K. N. D’Orazio, L. N. Lessen, R. Loll-Krippleber, G. W. Brown, R. Green, Distinct elongation stalls during translation are linked with distinct pathways for mRNA degradation. Elife 11 (2022).

120. C. C.-C. Wu, B. Zinshteyn, K. A. Wehner, R. Green, High-Resolution Ribosome Profiling Defines Discrete Ribosome Elongation States and Translational Regulation during Cellular Stress. Mol. Cell 73, 959–970.e5 (2019).

121. S. Souquere, S. Mollet, M. Kress, F. Dautry, G. Pierron, D. Weil, Unravelling the ultrastructure of stress granules and associated P-bodies in human cells. J. Cell Sci. 122, 3619–3626 (2009).

122. C. Zurla, A. W. Lifland, P. J. Santangelo, Characterizing mRNA interactions with RNA granules during translation initiation inhibition. PLoS One 6, e19727 (2011).

123. A. M. English, K. M. Green, S. L. Moon, A (dis)integrated stress response: Genetic diseases of eIF2α regulators. Wiley Interdiscip. Rev. RNA 13, e1689 (2022).

124. A. Eulalio, I. Behm-Ansmant, D. Schweizer, E. Izaurralde, P-body formation is a consequence, not the cause, of RNA-mediated gene silencing. Mol. Cell. Biol. 27, 3970–3981 (2007).

125. I. Horvathova, F. Voigt, A. V. Kotrys, Y. Zhan, C. G. Artus-Revel, J. Eglinger, M. B. Stadler, L. Giorgetti, J. A. Chao, The Dynamics of mRNA Turnover Revealed by Single-Molecule Imaging in Single Cells. Mol. Cell 68, 615–625.e9 (2017).

126. D. Zheng, N. Ezzeddine, C.-Y. A. Chen, W. Zhu, X. He, A.-B. Shyu, Deadenylation is prerequisite for P-body formation and mRNA decay in mammalian cells. J. Cell Biol. 182, 89–101 (2008).

127. L. A. Blake, L. Watkins, Y. Liu, T. Inoue, B. Wu, A rapid inducible RNA decay system reveals fast mRNA decay in P-bodies. Nat. Commun. 15, 2720 (2024).

128. G. P. Pijlman, A. Funk, N. Kondratieva, J. Leung, S. Torres, L. van der Aa, W. J. Liu, A. C. Palmenberg, P.-Y. Shi, R. A. Hall, A. A. Khromykh, A highly structured, nuclease-resistant, noncoding RNA produced by flaviviruses is required for pathogenicity. Cell Host Microbe 4, 579–591 (2008).

129. N. Kedersha, P. Ivanov, P. Anderson, Stress granules and cell signaling: more than just a passing phase? Trends Biochem. Sci. 38, 494–506 (2013).

130. K. Thedieck, B. Holzwarth, M. T. Prentzell, C. Boehlke, K. Kläsener, S. Ruf, A. G. Sonntag, L. Maerz, S.-N. Grellscheid, E. Kremmer, R. Nitschke, E. W. Kuehn, J. W. Jonker, A. K. Groen, M. Reth, M. N. Hall, R. Baumeister, Inhibition of mTORC1 by astrin and stress granules prevents apoptosis in cancer cells. Cell 154, 859–874 (2013).

131. F. Wippich, B. Bodenmiller, M. G. Trajkovska, S. Wanka, R. Aebersold, L. Pelkmans, Dual specificity kinase DYRK3 couples stress granule condensation/dissolution to mTORC1 signaling. Cell 152, 791–805 (2013).

132. P. Anderson, N. Kedersha, Stress granules: the Tao of RNA triage. Trends Biochem. Sci. 33, 141–150 (2008).

133. S.-H. Kim, S. Bae, M. Song, Recent development of aminoacyl-tRNA synthetase inhibitors for human diseases: A future perspective. Biomolecules 10, 1625 (2020).

134. G. H. M. Vondenhoff, A. Van Aerschot, Aminoacyl-tRNA synthetase inhibitors as potential antibiotics. Eur. J. Med. Chem. 46, 5227–5236 (2011).

135. F. L. Rock, W. Mao, A. Yaremchuk, M. Tukalo, T. Crépin, H. Zhou, Y.-K. Zhang, V. Hernandez, T. Akama, S. J. Baker, J. J. Plattner, L. Shapiro, S. A. Martinis, S. J. Benkovic, S. Cusack, M. R. K. Alley, An antifungal agent inhibits an aminoacyl-tRNA synthetase by trapping tRNA in the editing site. Science 316, 1759–1761 (2007).

136. E. M. Novoa, N. Camacho, A. Tor, B. Wilkinson, S. Moss, P. Marín-García, I. G. Azcárate, J. M. Bautista, A. C. Mirando, C. S. Francklyn, S. Varon, M. Royo, A. Cortés, L. Ribas de Pouplana, Analogs of natural aminoacyl-tRNA synthetase inhibitors clear malaria in vivo. Proc. Natl. Acad. Sci. U. S. A. 111, E5508–17 (2014).

137. M. Pines, D. Snyder, S. Yarkoni, A. Nagler, Halofuginone to treat fibrosis in chronic graft-versus-host disease and scleroderma. Biol. Blood Marrow Transplant. 9, 417–425 (2003).

138. M.-K. Park, J.-S. Park, E.-M. Park, M.-A. Lim, S.-M. Kim, D.-G. Lee, S.-Y. Baek, E.-J. Yang, J.-W. Woo, J. Lee, S.-K. Kwok, H.-Y. Kim, M.-L. Cho, S.-H. Park, Halofuginone ameliorates autoimmune arthritis in mice by regulating the balance between Th17 and Treg cells and inhibiting osteoclastogenesis: Halofuginone suppresses autoimmune arthritis. Arthritis Rheumatol. 66, 1195–1207 (2014).

139. L. L. de Figueiredo-Pontes, P. A. Assis, B. A. A. Santana-Lemos, R. H. Jácomo, A. S. G. Lima, A. B. Garcia, C. H. Thomé, A. G. Araújo, R. A. Panepucci, M. A. Zago, A. Nagler, R. P. Falcão, E. M. Rego, Halofuginone has anti-proliferative effects in acute promyelocytic leukemia by modulating the transforming growth factor beta signaling pathway. PLoS One 6, e26713 (2011).

140. J. Wang, L. Guan, J. Yu, B. Ma, H. Shen, G. Xing, Y. Xu, Q. Li, J. Liu, Q. Xu, W. Shi, J. He, Y. Huang, D. Yin, W. Li, R. Wang, Halofuginone prevents inflammation and proliferation of high-altitude pulmonary hypertension by inhibiting the TGF-β1/Smad signaling pathway. Sci. Rep. 15, 3619 (2025).

141. T. J. Carlson, A. Pellerin, I. M. Djuretic, C. Trivigno, S. B. Koralov, A. Rao, M. S. Sundrud, Halofuginone-induced amino acid starvation regulates Stat3-dependent Th17 effector function and reduces established autoimmune inflammation. J. Immunol. 192, 2167–2176 (2014).

142. Y. Hou, D. Wei, Z. Zhang, T. Lei, S. Li, J. Bao, H. Guo, L. Tan, X. Xie, Y. Zhuang, Z. Lu, Y. Zhao, Downregulation of nutrition sensor GCN2 in macrophages contributes to poor wound healing in diabetes. Cell Rep. 43, 113658 (2024).

143. S. A. Dogan, C. Pujol, P. Maiti, A. Kukat, S. Wang, S. Hermans, K. Senft, R. Wibom, E. I. Rugarli, A. Trifunovic, Tissue-specific loss of DARS2 activates stress responses independently of respiratory chain deficiency in the heart. Cell Metab. 19, 458–469 (2014).

144. T. Agnew, M. Goldsworthy, C. Aguilar, A. Morgan, M. Simon, H. Hilton, C. Esapa, Y. Wu, H. Cater, L. Bentley, C. Scudamore, J. Poulton, K. J. Morten, K. Thompson, L. He, S. D. M. Brown, R. W. Taylor, M. R. Bowl, R. D. Cox, A Wars2 mutant mouse model displays OXPHOS deficiencies and activation of tissue-specific stress response pathways. Cell Rep. 25, 3315–3328.e6 (2018).

145. T. Suzuki, The expanding world of tRNA modifications and their disease relevance. Nat. Rev. Mol. Cell Biol. 22, 375–392 (2021).

146. E. A. Orellana, E. Siegal, R. I. Gregory, tRNA dysregulation and disease. Nat. Rev. Genet. 23, 651–664 (2022).

147. T. Chujo, K. Tomizawa, Human transfer RNA modopathies: diseases caused by aberrations in transfer RNA modifications. FEBS J. 288, 7096–7122 (2021).

148. S. T. Gamage, R. Khoogar, S. H. Manage, M. C. Crawford, J. Georgeson, B. V. Polevoda, C. Sanders, K. A. Lee, K. D. Nance, V. Iyer, A. Kustanovich, M. Perez, C. T. Thu, S. R. Nance, R. Amin, C. N. Miller, R. J. Holewinski, T. Meyer, V. Koparde, A. Yang, P. Jailwala, J. T. Nguyen, T. Andresson, K. Hunter, S. Gu, B. A. Mock, E. F. Edmondson, S. Difilippantonio, R. Chari, S. Schwartz, M. R. O’Connell, C. Chih-Chien Wu, J. L. Meier, Transfer RNA acetylation regulates in vivo mammalian stress signaling. bioRxivorg, 2024.07.25.605208 (2024).

149. S. Blanco, S. Dietmann, J. V. Flores, S. Hussain, C. Kutter, P. Humphreys, M. Lukk, P. Lombard, L. Treps, M. Popis, S. Kellner, S. M. Hölter, L. Garrett, W. Wurst, L. Becker, T. Klopstock, H. Fuchs, V. Gailus-Durner, M. Hrabĕ de Angelis, R. T. Káradóttir, M. Helm, J. Ule, J. G. Gleeson, D. T. Odom, M. Frye, Aberrant methylation of tRNAs links cellular stress to neuro-developmental disorders. EMBO J. 33, 2020–2039 (2014).

150. J. E. Hurtig, A. van Hoof, An unknown essential function of tRNA splicing endonuclease is linked to the integrated stress response and intron debranching. Genetics 224, iyad044 (2023).

151. R. Moir, E. Merheb, V. Chitu, E. R. Stanley, I. M. Willis, Molecular basis of neurodegeneration in a mouse model of Polr3-related disease. Elife 13 (2024).

152. J. C. Darnell, S. J. Van Driesche, C. Zhang, K. Y. S. Hung, A. Mele, C. E. Fraser, E. F. Stone, C. Chen, J. J. Fak, S. W. Chi, D. D. Licatalosi, J. D. Richter, R. B. Darnell, FMRP stalls ribosomal translocation on mRNAs linked to synaptic function and autism. Cell 146, 247–261 (2011).

153. B. Popper, M. Bürkle, G. Ciccopiedi, M. Marchioretto, I. Forné, A. Imhof, T. Straub, G. Viero, M. Götz, R. Schieweck, Ribosome inactivation regulates translation elongation in neurons. J. Biol. Chem. 300, 105648 (2024).

154. J. J. Langille, K. Ginzberg, W. S. Sossin, Polysomes identified by live imaging of nascent peptides are stalled in hippocampal and cortical neurites. Learn. Mem. 26, 351–362 (2019).

155. M. N. Anadolu, J. Sun, S. Kailasam, K. Chalkiadaki, K. Krimbacher, J. T.-Y. Li, T. Markova, S. M. Jafarnejad, F. Lefebvre, J. Ortega, C. G. Gkogkas, W. S. Sossin, Ribosomes in RNA granules are stalled on mRNA sequences that are consensus sites for FMRP association. J. Neurosci. 43, 2440–2459 (2023).

156. Y. Kanai, N. Dohmae, N. Hirokawa, Kinesin transports RNA: isolation and characterization of an RNA-transporting granule. Neuron 43, 513–525 (2004).

157. R. El Fatimy, L. Davidovic, S. Tremblay, X. Jaglin, A. Dury, C. Robert, P. De Koninck, E. W. Khandjian, Tracking the Fragile X Mental Retardation Protein in a highly ordered neuronal RiboNucleoParticles population: A link between stalled polyribosomes and RNA granules. PLoS Genet. 12, e1006192 (2016).

158. G. Elvira, S. Wasiak, V. Blandford, X.-K. Tong, A. Serrano, X. Fan, M. del Rayo Sánchez-Carbente, F. Servant, A. W. Bell, D. Boismenu, J.-C. Lacaille, P. S. McPherson, L. DesGroseillers, W. S. Sossin, Characterization of an RNA granule from developing brain. Mol. Cell. Proteomics 5, 635–651 (2006).

159. E. Barbarese, D. E. Koppel, M. P. Deutscher, C. L. Smith, K. Ainger, F. Morgan, J. H. Carson, Protein translation components are colocalized in granules in oligodendrocytes. J. Cell Sci. 108 **( Pt** **8**), 2781–2790 (1995).

160. C. Müller, N. M. Bauer, I. Schäfer, R. White, Making myelin basic protein -from mRNA transport to localized translation. Front. Cell. Neurosci. 7, 169 (2013).

161. R. Smith, R. J. Rathod, S. Rajkumar, D. Kennedy, Nervous translation, do you get the message? A review of mRNPs, mRNA-protein interactions and translational control within cells of the nervous system. Cell. Mol. Life Sci. 71, 3917–3937 (2014).

162. K. C. Stein, J. Frydman, The stop-and-go traffic regulating protein biogenesis: How translation kinetics controls proteostasis. J. Biol. Chem. 294, 2076–2084 (2019).

163. M. A. Collart, B. Weiss, Ribosome pausing, a dangerous necessity for co-translational events. Nucleic Acids Res. 48, 1043–1055 (2020).

164. M. Aguilar Rangel, K. Stein, J. Frydman, A machine learning approach uncovers principles and determinants of eukaryotic ribosome pausing. Sci. Adv. 10, eado0738 (2024).

165. D. E. Weinberg, P. Shah, S. W. Eichhorn, J. A. Hussmann, J. B. Plotkin, D. P. Bartel, Improved ribosome-footprint and mRNA measurements provide insights into dynamics and regulation of yeast translation. Cell Rep. 14, 1787–1799 (2016).

166. T. Tuller, A. Carmi, K. Vestsigian, S. Navon, Y. Dorfan, J. Zaborske, T. Pan, O. Dahan, I. Furman, Y. Pilpel, An evolutionarily conserved mechanism for controlling the efficiency of protein translation. Cell 141, 344–354 (2010).

167. C. A. P. Joazeiro, Ribosomal stalling during translation: Providing substrates for ribosome-associated protein quality control. Annu. Rev. Cell Dev. Biol. 33, 343–368 (2017).

168. N. R. Guydosh, R. Green, Dom34 rescues ribosomes in 3’ untranslated regions. Cell 156, 950–962 (2014).

169. J. Schindelin, I. Arganda-Carreras, E. Frise, V. Kaynig, M. Longair, T. Pietzsch, S. Preibisch, C. Rueden, S. Saalfeld, B. Schmid, J.-Y. Tinevez, D. J. White, V. Hartenstein, K. Eliceiri, P. Tomancak, A. Cardona, Fiji: an open-source platform for biological-image analysis. Nat. Methods 9, 676–682 (2012).

