## Supplemental Figures for "tRNA synthetase activity is required for stress granule and P-body assembly"

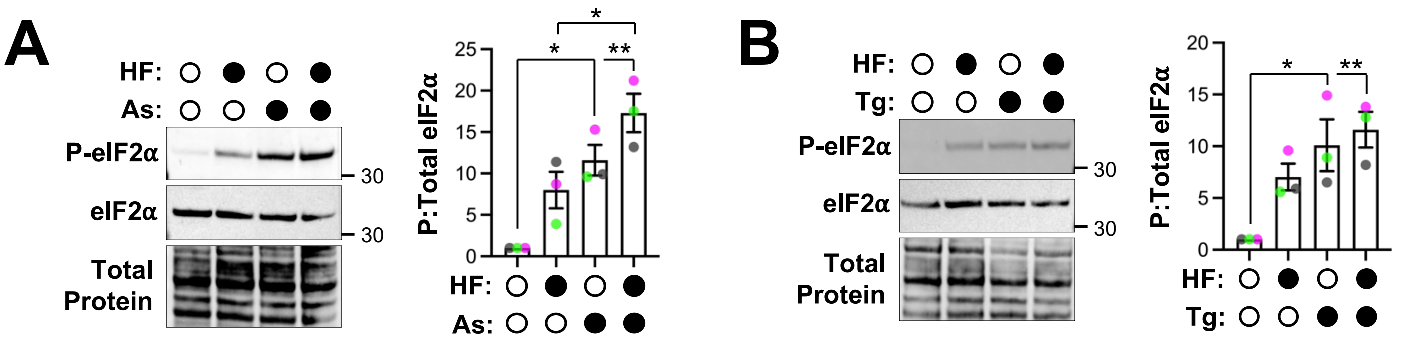


**Figure S1. Halofuginone co-treatment does not alter eIF2α phosphorylation by other stressors.** A. U-2 OS cells were treated with 0.2% DMSO carrier, 20 μM halofuginone (HF), 250 μM arsenite (As), or both HF and As for 1 hour, followed by western blotting for phosphorylated and total eIF2α with total protein shown. The ratio of P-eIF2α:eIF2α is quantified from n=3 independent experiments. Cells were treated with 0.2% DMSO, 20 μM HF, 1.25 μM thapsigargin (Tg), or both HF and Tg for 1 hour and western blotting done as in (A) from n=3 independent experiments. Representative images shown with average +/- s.e.m. reported, with each dot representing the average of one independent replicate (in gray, green, or pink). Molecular weights (kDa) indicated on each blot. Statistical significance was assessed with an ordinary one-way ANOVA followed by Tukey's multiple comparisons test with * $p \leq 0.05$ and ** $p \leq 0.01$.

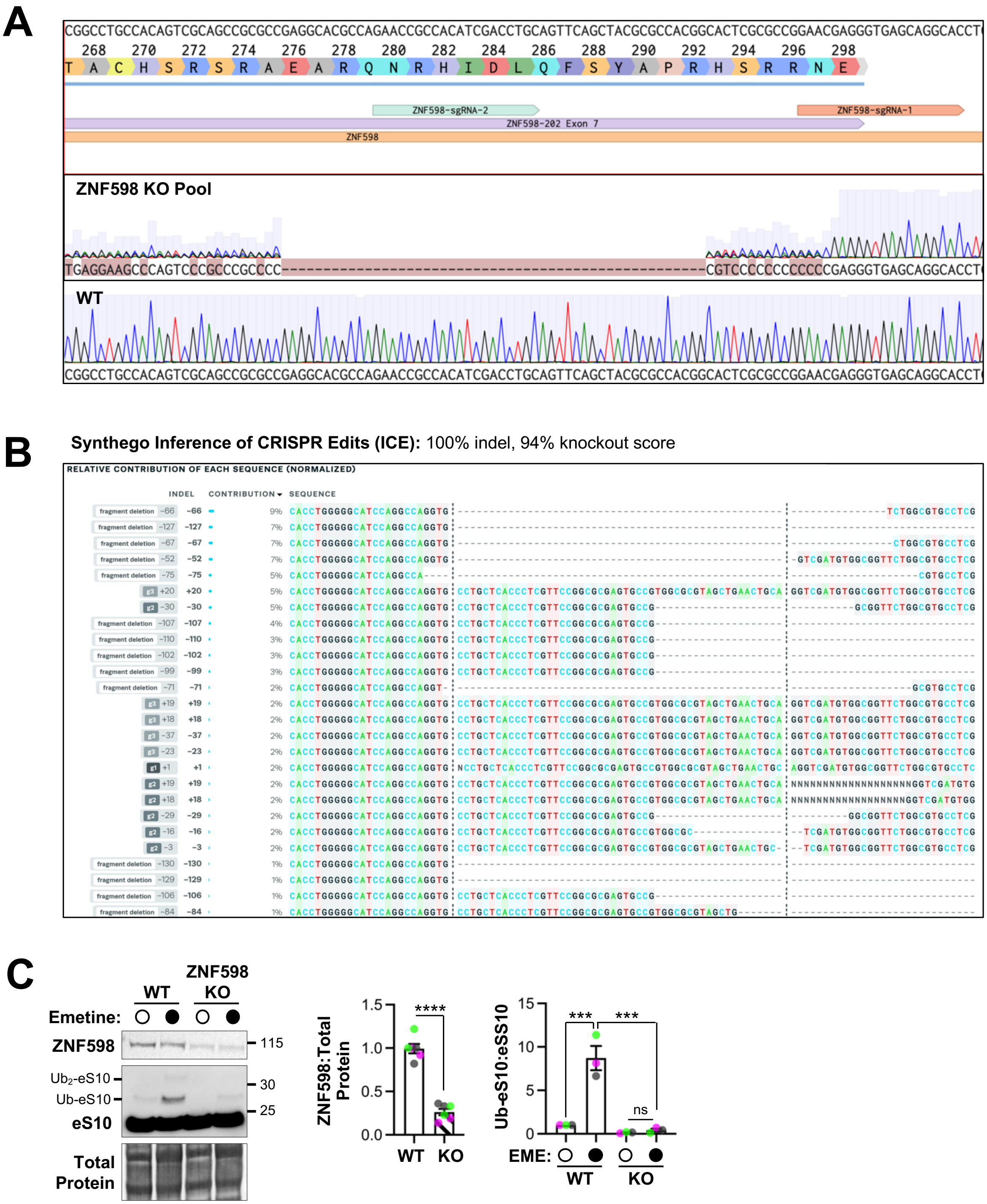


**Figure S2. Validation of ZNF598 knockout in U-2 OS cells.** A. Alignment of Sanger sequenced PCR amplicons from WT GFP-G3BP1 U-2 OS versus those from the ZNF598 CRISPR-Cas9 knockout (KO) cell pool. Annotations show placement of two of three gRNAs that were used to create large deletions in the ZNF598 coding region. B. Results of Interference of CRISPR Edits (ICE) on Sanger sequencing data from (A). C. Western blotting of WT and ZNF598 KO GFP-G3BP1 U-2 OS cells treated with either DMSO carrier (0.2%) or emetine (0.18 µM) for 15 minutes of ZNF598, eS10, with total protein shown. ZNF598 levels relative to total protein were determined and Ub-eS10:eS10 quantified and reported at right, with a representative blot at left. The average +/- s.e.m. from n=3 independent experiments is shown with independent replicates indicated as pink, gray, or green dots. Molecular weights (kDa) indicated on each blot. Significance for differences in ZNF598 protein levels between WT and KO was assessed with Student's t-test (**** $p \leq 0.0001$), and ordinary one-way ANOVA with Tukey's multiple comparisons test done for the Ub-eS10:eS10 ratios (*** $p \leq 0.005$, ns = not significant).

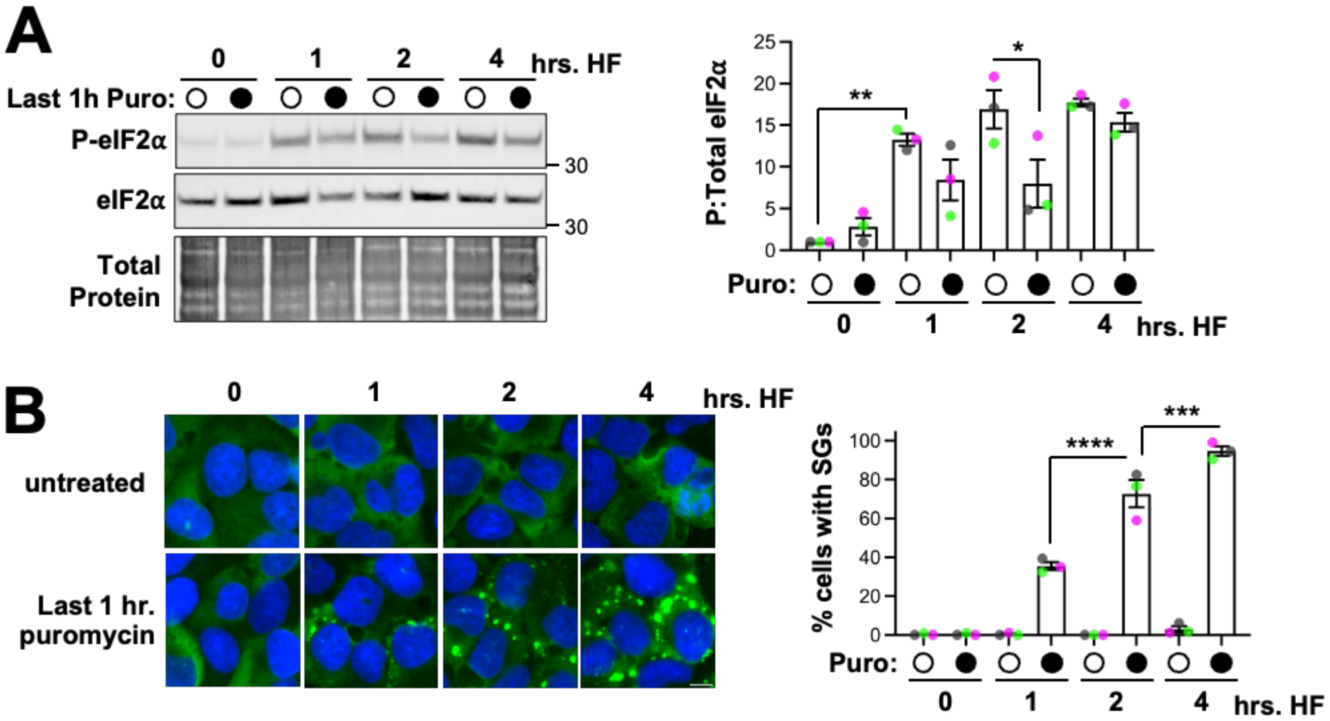


**Figure S3. The effects of puromycin on P-eIF2α and SG assembly over time during halofuginone treatment.** A. U-2 OS cells expressing GFP-G3BP1 were treated with HF (20 µM) for 1, 2, or 4 hours in the presence or absence of puromycin (10 µg/mL) added 1 hour prior to collection. Western blotting and quantification of phosphorylated and total eIF2α was done from n=3 independent experiments (representative blot shown at left and average +/- s.e.m. shown at right). B. Cells were treated as in (A) and SGs were detected and quantified from n=3 independent experiments, with ≥292 cells counted per treatment across all replicates. Scale bars are 10 µm. Quantification is reported as mean with s.e.m. shown, with gray, green, and pink dots indicating each replicate average. Molecular weights (kDa) indicated on each blot. Statistical significance was assessed with an ordinary one-way ANOVA followed by Tukey's multiple comparisons test: $^*p \leq 0.05$, $^{**} p \leq 0.01$, $^{***} p \leq 0.005$, $^{****} p \leq 0.001$.

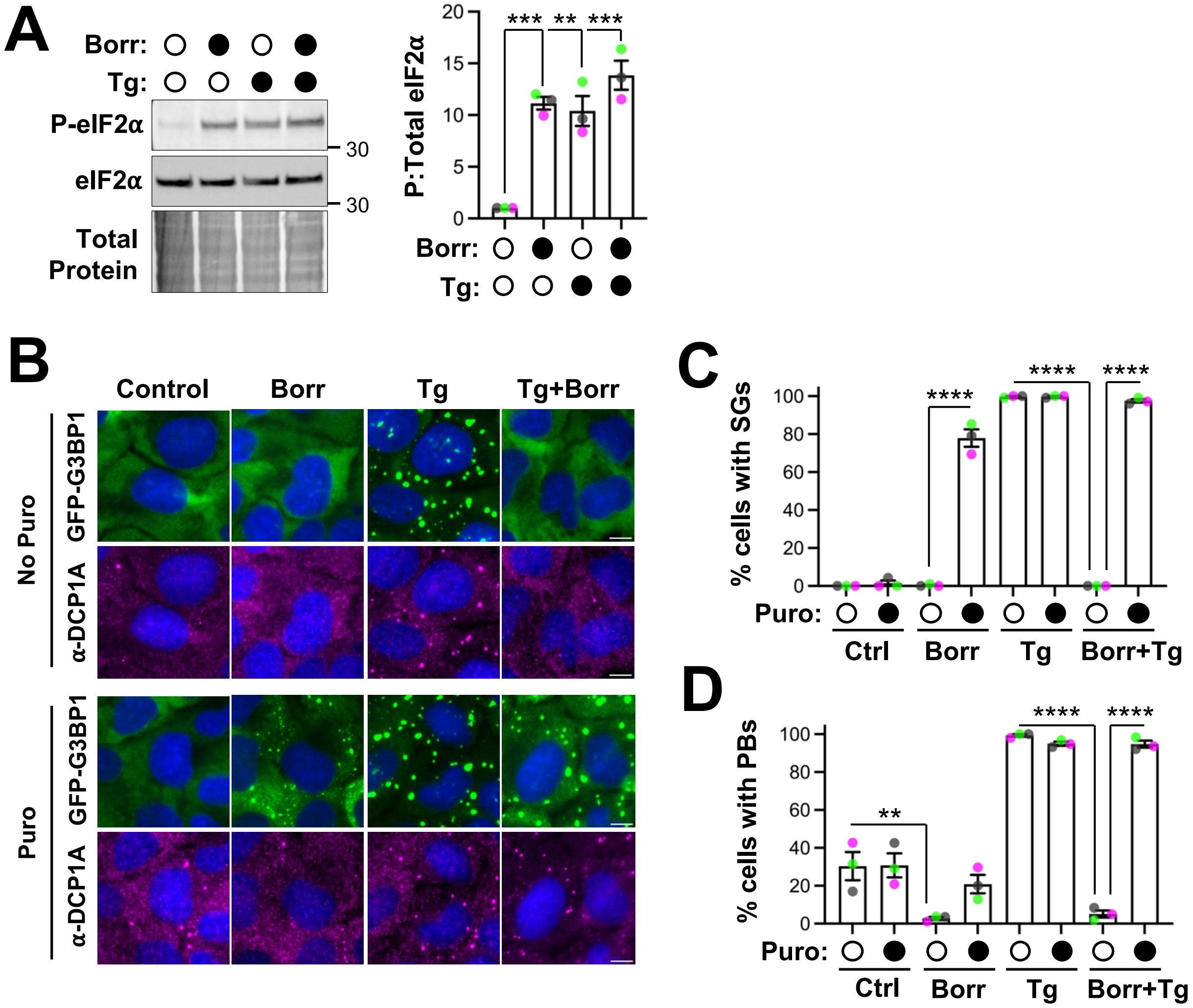


**Figure S4. Borrelidin induces the ISR and inhibits RNP granule assembly in a manner rescued by puromycin.** A. U-2 OS cells expressing GFP-G3BP1 were treated with either borrelidin (Borr, 100 µM) or ethanol (1%) as carrier control for 3 hours, with or without thapsigargin (Tg, 1.25 µM) or DMSO added 1 hour prior to collection. Western blotting for phosphorylated and total eIF2α was performed, and representative blots shown with total protein at left; at right are average +/- s.e.m. from three independent experiments. Molecular weights (kDa) shown on each blot. B. Cells were treated as in (A) in the presence or absence of puromycin (10 µg/mL) added 1 hour prior to fixation. Immunofluorescence imaging was performed for P body (PB) marker DCP1A alongside imaging of the GFP-G3BP1 stress granule (SG) marker. Shown are representative images from n = 3 independent experiments. The percent cells with SGs (C) and the percent cells with PBs (D) is shown with ≥307 cells counted per treatment across all replicates, respectively. The averages +/- s.e.m. are shown from n=3 independent experiments. All scale bars are 10 µm. Significance was assessed with ordinary one-way ANOVAs and Tukey's multiple comparisons tests with ** p. ≤0.01, *** p≤0.005, **** p≤0.001.
